# Structural and Metabolic Characterization of Ni(I)-inhibitors Provide a Robust Anti-Methanogenicity Scoring System

**DOI:** 10.64898/2026.02.25.708075

**Authors:** Randy Aryee, Mohammad Reza Zargar, SR Vaishnavey, B. Arunraj, Noor S. Mohammed, Supantha Dey, Karuna Anna Sajeevan, A. Nathan Frazier, Jacek A. Koziel, Matthew R. Beck, Thomas J. Mansell, Ratul Chowdhury

**Affiliations:** Department of Chemical and Biological Engineering, Iowa State University, Ames, Iowa, USA; The Center for Biorenewable Chemicals, Iowa State University, Ames, Iowa, USA; Maseeh Department of Civil, Architectural and Environmental Engineering, University of Texas, Austin, Texas, USA; USDA-ARS Conservation and Production Research Laboratory, Bushland, Texas, USA; Department of Animal Science, Texas A&M University, College Station, Texas, USA

**Keywords:** enteric methanogenesis, methyl coenzyme reductase, enzyme inhibition, methane mitigation, contrastive machine learning, metabolic modeling

## Abstract

Atmospheric methane (CH4) acts as a key contributor to global warming and a short-lived climate forcer. CH4 mitigation represents the most promising means to address short-term climate change. Ruminant enteric CH4 produced by methanogenic archaea represents 27.2% of global CH4 emissions. Only a few of the direct methanogenesis inhibitors identified bear high mitigation potential hence it is important to investigate their underlying modes of action. Here, we elucidated biophysical and thermodynamic interplay between known inhibitors and cofactor F430, to determine their stoichiometric ratios and binding affinities. We leverage this prior in a robust contrastive learning approach to functionally cluster known sixteen inhibitors and 53,959 bovine-linked metabolites. We demonstrate a multi-factor optimization protocol to identify putative inhibitors with: (i) high bacterial membrane permeability, (ii) no adverse effect to ruminal fermentation, (iii) known degradation pathway, and (iv) direct commercial availability. Subsequent *in vitro* assays and community metabolic modeling with a first set of eight treatment molecules revealed structo-metabolic priors that tie thermodynamic signatures of inhibition to metabolic flux shifts. We established a multi-scale workflow that transforms ostensibly negative compounds into mechanistic insight, linking rumen metabolic flux shifts to MCR–F430-Ni(I) inhibition chemistry as a foundation for rational methane-mitigation design.

**COVER ABSTRACT:** 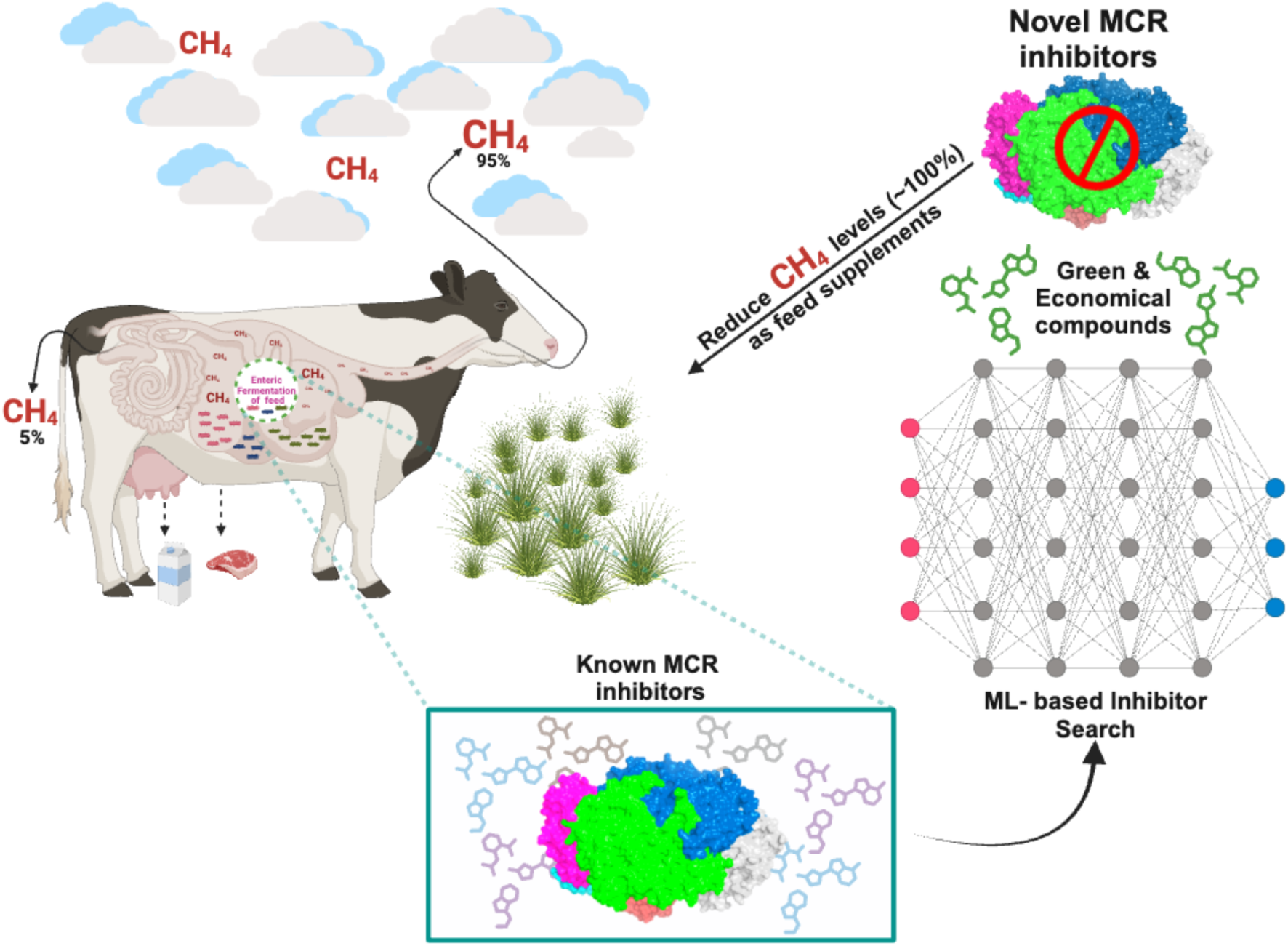

## 1. Introduction

Greenhouse gases (GHGs) possess the potential to absorb and re-radiate infrared radiation in the atmosphere. This ability initiates the trapping of heat, resulting in a rise of global surface temperature, a means that leads to global warming over long periods of occurrence^1,2^. Prominent GHGs include carbon dioxide (CO_2_), methane (CH_4_), nitrous oxide (N_2_O), and a hand full of fluorinated gases^1–4^. GHGs have extensively been one of the world’s major climate change drivers over generations. Inclusive of key anthropogenic activities leading to global warming are enteric CH_4_ emissions from ruminant livestock, the release of CO_2_ from fossil fuel use, land use change, and landfills.

According to the sixth assessment report by the Intergovernmental Panel on Climate Change (IPCC), there were 59 Gt of CO_2_-equivalence (CO_2_-e) emitted globally in 2019^5^. Emissions from Agriculture, Forestry and Other Land Use (AFOLU) represented 22% of these emissions. Enteric CH_4_ emissions accounted for 5.1% of total global GHGs, 23% of AFOLU, and 27.2% of total CH_4_ emissions **(Figure 1)**. As CH_4_ has a short atmospheric lifespan (approximately 12 yr), in periods where emission rates are reduced to a large enough degree, there will be less atmospheric CH_4_, resulting in lower warming^6^. Accordingly, models predict that rapidly declining CH_4_ emissions can reduce temperature equivalent to the removal of atmospheric CO_2_^5^. As such, reductions in CH_4_ emissions represent the most promising means to address climate change in the short term^7^. This nuance of CH_4_ emissions in general and the relative contribution of enteric CH_4_ to total CH_4_ emissions make enteric CH_4_ mitigation particularly important^6,8^.

**Figure 1.**
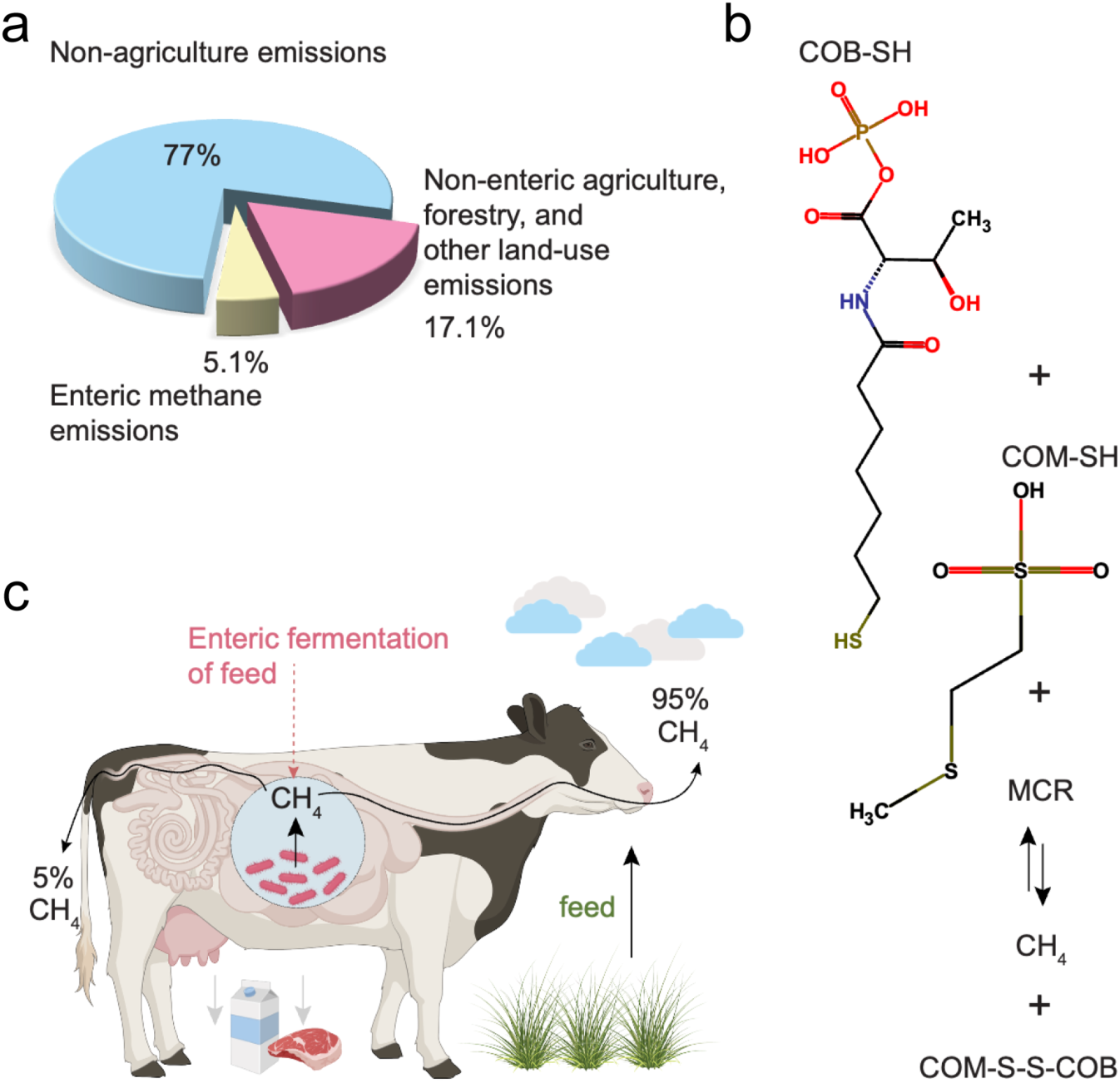
Representative schematic illustrating the distribution of greenhouse gas emissions, focusing specifically on CH_4_ and detailing its biochemical synthesis and release mechanisms. **a)** Global representation of GHGs emissions with distributions centered on CH_4_ by sector as gathered from literature **b)** The entire enteric fermentation of carbohydrate (cellulose) feed as a mechanism of CH_4_ release. **c)** Biochemical reaction and the rate-limiting step in enteric CH_4_ synthesis catalyzed by MCR enzyme.

Due to increasing production efficiency, the carbon footprint of beef production in the U.S. was reduced by 16.3%^9^. Similarly, the carbon footprint of dairy production in the U.S. was reduced by 19.3% from 2007 to 2017, from 1.66 to 1.34 kg of CO_2_-e per kg of milk^9^. While these reductions in carbon footprint from the dairy and beef industry are commendable, recent pledges of carbon neutrality by industries and companies have increased since the Paris Climate Agreement^10^. These types of commitments require reductions in absolute emissions rather than reductions in emissions per unit of product. Accordingly, enteric CH_4_ mitigation is highly needed by both the dairy and beef industries.

Methanogenesis, in brief, defines CH_4_ biosynthesis, irrespective of its emission source. Methane is a key natural secondary metabolite of enteric fermentation in the rumen of ruminants upon the digestion of consumed feed^11^. By utilizing hydrogen, methanogenesis provides an important role in maintaining homeostasis in the ruminal environment. Methanogens (methanogenic archaea) are the predominant mediators of methanogenesis within the rumen^12^. By impacting the availability of substrate, methanogens and CH_4_ production are greatly influenced by other microbial members, primarily bacteria^13^. Accordingly, the majority of enteric CH_4_ mitigation strategies, including alternative hydrogen sinks, shifts in fermentation parameters, and feed management impact substrate availability and subsequently decrease CH_4_^12^. However, methanogens represent a key target for investigating metabolic processes for CH_4_ mitigation, with direct methanogenesis inhibitors representing the largest mitigation potential^14,15^

Significantly, enteric CH_4_ production has been a conventional marker for farming productivity as CH_4_ is an associated product for carbohydrate utilization in ruminants. Also, the quest for essential and volatile fatty acid production in livestock dietary metabolism has leveraged this gross implication of CH_4_ production in the four-chambered stomach of herbivorous grazing mammals^16^. Again, a natural result of excess hydrogen eradication in ruminants leads to CH_4_ being released into the atmosphere through either eructation (95%) or flatulence (5%)^1^ (**Figure 1**). Following a stepwise biochemical reaction of CH_4_ biogenesis in ruminants, the enzyme Methyl Coenzyme M Reductase (MCR) produced from methanogenic archaea plays a key role ^1,2,17^. MCR catalyzes the final but rate-limiting step between methyl-coenzyme B (COB-HS) and methyl-coenzyme M (CH_3_-S-COM) to release a heterodisulfide Coenzyme M and Coenzyme B (COM-S-S-COB) and CH_4_ as products^18^ (**Figure 1**). The entire biochemical process is labeled methanogenesis for reference^19^.

Biological methane mitigation will not be unlocked by a single *glory* molecule discovered through a single round of metric-driven screening. What is needed instead is an interpretable, structure-to-metabolism framework that treats even sub-optimal compounds as mechanistic probes—capable of revealing how rumen fluxes, digestion efficiency, and MCR–F_430_-Ni(I) chemistry are coupled across scales. This is the key scientific motivation behind this endeavor. Methanogenesis mitigation strategies and approaches are being actively conceptualized, designed, and deployed for a green and CH_4_-reduced ecosystem. Currently explored CH_4_ mitigation strategies include promising options such as increasing feeding levels, decreasing dietary forage-to-concentrate ratios, improving feed quality and digestibility. However, these strategies often reduce enteric CH_4_ emission on a per product produced basis and have only demonstrated reductions by around 16.3%^15^. Mitigation options that reduce absolute emissions have also been investigated and include genetic and breeding selection^20^, feeding tanniferous forages^21^, providing electron sinks^22^, and supplementing fat^23,24^. These options have shown to reduce enteric CH_4_ emissions by around 10%^15^.

The mitigation options that have demonstrated the largest enteric CH_4_ mitigation potential are direct methanogenesis inhibitors. These include 3-nitrooxypropanol (3-NOP) and bromoform (CHBr_3_)-containing seaweeds *(Asparagopsis* spp.). 3-NOP has been shown to reduce enteric CH_4_ by 25-30% ^25^ and *Asparagopsis* seaweeds have reduced enteric CH_4_ by 80-98% ^26,27^. 3-NOP and CHBr_3_ from red seaweed inhibit methanogenesis by competitively binding and providing an agonistic effect on CH_3_-S-COM, hence hindering the final and rate-limiting step in enteric methanogenesis^1,28,29^. More specifically, halogenated compounds such as CHBr_3_ competitively displace other natural substrates that tend to interact with the Ni(I) ion of F_430_ coenzyme M. This results in methyl transfer inhibition and a reduction in CH_3_-S-COM mediated CH_4_ release^29^. 3-NOP, while promising, have demonstrated very low adoption rates and preclude new intellectual property and commercialization of its intuitive derivatives (like 3-diglyceryl nitrate, n-propyl nitrate) for commercial scale up without infringing 3-NOP’s patent guardrails. This work adheres to this premise and uncovers a list of structural, biochemical, metabolic and commercial scoring metrics that can expedite discovery of novel, commercializable anti-methanogenic molecules.

Amongst the methanogenesis inhibitors investigated so far, a data-driven deep-dive with precise molecular multi-scale modeling of the atomic-level Ni(I)-biochemistry to community-level metabolism of these inhibition mechanisms along with *in vitro* testing that targets maximizing total gas while minimizing methane concentration has remained largely elusive. Empirical approaches by far have not provided enough biochemical information on Ni(I) chemistry to identify or design novel inhibitor molecules which have a high binding affinity to rumen MCR’s cognate cofactor Ni(I)-F_430_, aside from 3-NOP^1,2^. Here, we employ detailed structural biology investigation to interpret the stoichiometric ratios (i.e., biophysical flooding), binding affinities (i.e., biochemical trapping), bacterial membrane permeability, and metabolic shifts induced by all 16 well-documented inhibitor molecules against the redox-active nickel (Ni(I)) complexed with tetrahydrocorphin coenzyme F_430_ cofactor of MCR. We further set up a contrastive machine learning (ML) approach to test a new generation of molecules for their anti-methanogenicity, that are also bovine-linked metabolites (i.e., accounted for downstream human food safety, and degradation or export pathways). We *in vitro* tested two rounds of four molecules in bovine ruminal fluid and catalog their anti-methanogenic potentials and volatile effluents.

## 2. Methods

### Structural biology of MCR dictate binding rules for competitive inhibitor compounds

A data mining sweep through reported literature was performed encompassing Web of Science, GenBank^30^, and UniProt^31^ databases to catalog the methanogenic archaea MCR enzyme responsible for enteric CH_4_ biosynthesis. Studies focused on the biochemical mechanism of the MCR enzyme, inhibitor molecules, and structural insights of both MCR and the inhibitor molecules were shortlisted. Based on the structural insight, a high-resolution, X-ray diffracted crystallized MCR (PDB Accession ID: 5G0R) was identified^2^ from the Research Collaboratory for Structural Bioinformatics Protein Data Bank (RCSB PDB)^32^. Protein structure visualization, characterization, and determination of active site residues within a 5 Å distance from the cofactor F_430_ were investigated using PyMOL^33,34^. A focused library of known anti-methanogenic small molecules (AMs) deep-mined from PubChem^35^ annotated with direct/ indirect experimental evidence was collated to this end. The MolView server^36^ was used to generate structures for inhibitors that were not available in PubChem.

### 2.1. Thermodynamics of docking of known anti-methanogenic molecules (AMs)

Molecular docking simulations were conducted to provide insights into the (a) non-clashing binding poses (i.e., flooding stoichiometries), and (b) thermodynamics of methanogenesis inhibition across known inhibitor molecules with the Ni(I) of F_430_ cofactor of MCR. Rigid molecular docking was performed using AutoDock Vina^37^ to explore the binding interactions of the selected inhibitors out of a library of literature-derived small molecule compounds and the cofactor F_430_ of MCR (PDB ID: 5G0R). Protein and ligand preparation steps were conducted using AutoDockTools^37^ and OpenBabel^38^. Configuration file for docking had fixed grid box values of x, y and z coordinates across all ligands. Using the gradient-based local search genetic algorithm built in AutoDock Vina ^37^, the docking energy scores and rankings of binding poses of each inhibitor molecule to the active-site of the MCR enzyme were obtained. Illustrations of inhibitor-MCR complex were generated using PyMOL^33,34^. Additionally, we performed Boltz-2^39^ template docking of the inhibitor compounds with the apo-MCR structure to rank complexes based on binding probability predictions. This was repeated for the predicted and tested molecules in this study.

### 2.2. Stoichiometric ratio and binding affinity analysis with MCR

The stoichiometric ratio and distribution of inhibitor molecules within the catalytic groove of MCR with cofactor F_430_ were analyzed using PyMOL. Inhibitors were profiled by first following the flooding rule within the MCR-F_430_ pocket (i.e., how many molecules can fit into the active site without self-clashing). All ligands’ poses within <5 Å from Ni(I) of cofactor F_430_ were classified into groups of two based on their site of binding. The number of such poses for each inhibitor represents the maximum biophysically permissible stoichiometric ratio against inhibitor molecule type.

### 2.3. Structural comparison of MCR inhibitors with ruminant specific metabolite databases

Sixteen known inhibitors explored against MCR enzyme were compared for similarities in molecular fragments within two ruminant specific metabolite databases - a) Milk Composition Database (MCDB) ^40^ and b) Bovine Metabolome Database (BMDB) ^41^, containing 2,322 and 51,637 entries, respectively. The structural information of metabolites was downloaded in Structure-Data File (SDF) format and further processed to obtain canonical Simplified Molecular Input Line Entry System (SMILES) representation using RDKit^42^. Further data curation process involved removing invalid SMILES, duplicate entries, and entries containing metal ions to ensure the data quality and relevance. The curated SMILES strings were then embedded using the ChemBERTa ^43^ model, a pre-trained model specifically trained on a dataset of 77 million molecule SMILES strings from PubChem^44^. The resulting 768-dimensional feature vectors represent each molecule used in the subsequent analysis. For both datasets, cosine similarity was calculated between the embeddings of the known anti-methanogenesis molecules and the metabolites. Bovine and milk metabolites with a cosine similarity of 85% or higher to the known anti-methanogenesis molecules (highly similar feature vectors) were selected and combined with the known anti-methanogenesis molecules to form a new dataset for contrastive learning. The new dataset comprising of metabolites with greater than or equal to 85% similarity were treated as the positive class, while the remaining metabolites from each dataset (bovine and milk) served as the negative class.

A contrastive learning approach was employed using triplet loss to refine the embedding space and enhance the separation between the positive and negative classes^45^. The triplet loss function minimized the distance between anchor-positive pairs while maximizing the distance between anchor-negative pairs, with a margin of 0.5^45,46^. The neural network architecture used for feature transformation was a feedforward network consisting of three hidden layers, followed by rectified linear unit (ReLU) activations, and an output layer projecting the embeddings into a 256-dimensional space. The training data was composed of triplets, where the anchor and positive samples came from the combined dataset (either known anti-methanogenesis molecules + ≥ 85%-similar milk metabolites or known anti-methanogenesis molecules + ≥ 85%-similar bovine metabolites), and the negative samples were selected from the remaining metabolites (milk or bovine). The model was trained for 1,300 epochs using the Adam optimizer with a learning rate of 1×10^-6^. During each epoch, triplets were passed through the network to compute embeddings, and the triplet loss was calculated and minimized through backpropagation. The final embeddings were visualized using t-Distributed Stochastic Neighbor Embedding (t-SNE) to validate the clustering, with similar molecules forming tight clusters in the reduced-dimensional space. All computations were performed using Python with PyTorch for deep learning, NumPy for numerical operations, and Matplotlib for visualization, utilizing a GPU-enabled environment to accelerate the training process.

### 2.4. Scoring molecules based on binding affinity and stability of selected Ni(I) inhibitor complexes

All potential inhibitor compounds from MCDB, BMDB and a well curated commercial Sigma-Aldrich (ease of market availability) dataset extracted from PubChem were investigated for chemical similarity to 3-NOP as prior for commercially available alternated of 3-NOP. For potential inhibitor compounds with more than 50% Tanimoto similarity to 3-NOP, we obtained binding likelihood and half-maximal inhibitory concentration (IC_50_)-like interaction predictions using Boltz-2^47^. To enhance the accuracy of the structure prediction module and confidence of binding affinity estimates, the macromolecule crystallographic information framework (mmCIF) structure of the receptor (PDB: 5G0R) was employed as a template. The input included the receptor sequence comprising chains A and D of MCR and SMILES representations of candidate inhibitors. All output complexes are ensured to have high interface predicted TM-score (ipTM) (>0.85) scores indicating reliable inter-chain packing and structural integrity. Trained on experimental and molecular dynamics ensembles, the affinity predictions are based on molecular conformers at the binding pocket and do not explicitly incorporate cofactor information. Quantitative affinity readouts were normalized to a continuous log_10_ scale (µM units) to allow relative ranking of inhibitor compounds. The binding stability constants (logK) of these selected compounds with Ni(I) in cofactor F430 are estimated using LOGKPREDICT v2.0.0^48,49^ and ChemProp v1.6.1 using the *chemprop_predict* module. LOGKPREDICT employs a directed message passing neural network (D-MPNN) on atomic feature-annotated molecular graphs of the metal-ligand complexes available in IUPAC and NIST databases to predict log K of binding between a metal and a chelator.

### 2.5. Quantifying toxicity of top 2% bovine linked metabolites that bear potential for MCR-binding

To score the toxicity of the 1,111 (top 2% ranked for MCR-binding,) bovine linked molecules, we leveraged the MolToxPred model ^50^. Each molecule was represented through molecular descriptors and Morgan fingerprints, which captured key structural and physicochemical properties. The model then analyzed these features and assigned, for each molecule, a toxicity score between 0 and 1, with 0 indicating lower toxicity and 1 representing higher toxicity. The top fifteen low-toxicity (<0.25) molecules were grouped into four categories based on their toxicity scores, permeability scores and similarity to known anti-methanogenic molecules, as measured by Euclidean distance. It must be noted that such toxicity predictors at best reflect the relative toxicity trends at some arbitrarily normalized concentrations. Since concentrations and buffer pH (i.e., dose-response curve data is scarce) do not factor into MolToxPred predictions, it remains blind to concentrations. For our *in vitro* experiments we used high dilutions of these feed additives, we assume this 0→1 predictions to be reasonable. A follow-up work will provide a rumen-specific, concentration-aware toxicity predictor. Additionally, to validate the top 15 predicted anti-methanogenic molecules from our contrastive learning model, we used the same docking approach as that of the 16 tested inhibitor-MCR complexes using AutoDock Vina^37^ the to obtain docking scores for the predicted molecules . Following the same restrained protein-ligand docking approach applied to known methanogenic molecules, we focused on predicted molecules that were positioned within 5 Å of the Ni(I) center of the F430 cofactor in MCR and determined their site of binding.

### 2.6. Quantifying a bacterial membrane permeability proxy score for molecules

We quantified the permeability (log Papp values) of selected molecules using an *in silico* Caco-2 permeability model from Falcón-Cano et al.^51^. The updated and implemented workflow involved cleaning and standardizing chemical structures, removing duplicates, and segregating measurements into a high-confidence set with standard deviation ≤ 0.5 and another set for recursive refinement. From 2D RDKit-derived physicochemical and shape descriptors plus Morgan fingerprints (1024 bits), a recursive random forest algorithm identified the five most predictive features: slogP, TPSA, SMR, Hall–Kier Alpha, and Kappa 3. We generated permeability predictions using their conditional consensus random forest architecture, which comprises one global model and four regional models stratified by permeability range. However, as Caco–2 monolayers model human intestinal epithelium rather than bacterial envelopes, these values should be considered a naive proxy for bacterial membrane permeability rather than a mechanistic description of bacterial uptake^52^.

### 2.7. In vitro testing of MCR-binding, non-toxic, high-permeability molecules in bovine rumen fluid

Initially, our framework identified imidazole, L-carnitine, methyl jasmonate and propylpyrazine as promising candidate molecules to be blind-tested MCR inhibition (before metabolic model integration). These compounds were purchased from Millipore Sigma for the *in vitro* screening experiment. The dose employed within this experiment was scaled per the reported optimal dose of 3-NOP of 60 mg/kg dry matter (DM). We normalized the dose of these compounds for molecular weight (MW of 3-NOP = 121.09 g/mol) and then multiplied this by the modeled stoichiometric ratio. Accordingly, the dosage (in mg/kg DM) used in the *in vitro* gas production screening was 101.2 for imidazole, 79.9 for L-carnitine, 111.1 for methyl jasmonate, and 60.5 for propylpyrazine.

All *in vitro* incubations were conducted using the ANKOM RF gas production system (ANKOM Technology, Macedon, NY). The system is composed of 250-mL jars (the incubation vessels) and a module, which transmits gas pressure data to a nearby analysis machine. The system was programed to record gas pressure every 10 min for the duration of the incubations (48-h). Cumulative gas volume (mL) was calculated from gas pressure by first calculating gas moles using the ideal gas law and then volume based on Avogadro’s Law, which states that 1 mol of gas will occupy 22.4 L at standard temperature and pressure. A control (no-added candidate molecules) and the 4 candidate methanogenesis inhibitors were replicated 3 times per incubation run (48-h) and the experiment consisted of 3 runs. This results in 9 replicates per treatment. During each run, 1-g of substrate was provided, and the dose of the candidate compounds were applied to their respective incubation vessels at the previously mentioned inclusion level. Each incubation vessel also received 80 mL of a buffer solution^53^ and 20 mL of freshly collected ruminal fluid. The ruminal fluid was obtained from 2 cannulated steers per incubation period (1-L from each steer), was strained through 4 layers of cheese cloth, and placed in pre-warmed (39 °C) thermoses. The fluid was then thoroughly mixed and added to the incubation vessels. Care was taken throughout the collection and application of ruminal fluid to the incubation vessels to maintain an anaerobic environment, with the headspace of thermoses, containers and incubation vessels purged with CO_2_. Two F57 bags (ANKOM Technology, Macedon, NY) were also added to each incubation vessel with 0.5 g of substrate contained within the filter bag. Clips (6.3 ± 0.30 g; mean ± SD) were attached to the filter bags to keep the bags submerged within the ruminal fluid and buffer. The diet fed to the cannulated steers and the substrate used were the same (**Table 1**). Each *in vitro* run also had blank jars, which contained only ruminal fluid and buffer solution and gas volume was adjusted based on the gas produced from these blanks.

**Table 1.** Composition of diets fed to cannulated steers and used as substrate in the *in vitro* experiments.

| Ingredient | Inclusion, % of DM |
| --- | --- |
| Wheat hay | 35 |
| Steam Flaked Corn | 35 |
| Wet corn gluten feed<sup>1</sup> | 20 |
| Dried distillers' grain plus solubles | 7 |
| Premix<sup>2</sup> | 3 |

<sup>1</sup> SweetBran (Cargill Sweet Bran, Dalhart, TX)
<sup>2</sup> Containing (DM basis): 27.3610% calcium carbonate, 22.6% dried distillers' grains plus solubles, 20.7% magnesium sulfate, 17.3% monocalcium phosphate, 7.02% salt, 4.03% potassium chloride, 0.34% Rumensin 90 (Elanco Animal Health; Greenfield, IN), 0.25% manganese sulfate, 0.14% vitamin E (500 IU/g), 0.1382% zinc oxide, 0.0798% copper sulfate, 0.043% sodium selenite, 0.0056% vitamin A (1,000,000 IU/g), 0.0015% cobalt carbonate, 0.0012% ethylenediamine dihydroiodide, 0.0011% vitamin D (500,000 IU/g).

After F57 bags, buffer, and ruminal fluid was added, initial pH of each jar was measured using a Laqua F-73G pH meter (Horiba Scientific; Kyoto, Japan). Finally, the headspace of each incubation vessel was purged with CO_2_ and then sealed with the ANKOM RF gas production modules. The vessels were then placed into a shaking incubator (Innova S44i, Eppendorf, Enfield, CT) at 39 °C and 50 rpm for 48 h. Throughout the incubation, vented gas was collected into sample bags (FlexFoil^®^ PLUS; SKC Ltd.; Eighty Four, PA). These gas samples were analyzed for CH_4_ concentration using an 8610C gas chromatograph (GC; SRI Instruments; Torrance, CA). Methane concentration and 48-h cumulative gas volume were multiplied to determine CH_4_ production (mL/g of DM). Additionally, following the 48-h incubations, the F57 bags were rinsed until the rinse water ran clear and dried in a forced air oven (55 °C) for 72-h. The F57 bags then underwent the neutral detergent fiber procedure without sodium sulfite or α-amylase in the ANKOM Delta Fiber Analyzer for the determination of *in vitro* true DM digestibility (IVTDMD)^54^.

Following the incubation period, vessels were removed from the incubator, final pH was measured (Laqua F-73G pH meter, Horiba Scientific, Kyoto, Japan), and ruminal fluid was subsampled into two, 2-mL microcentrifuge tubes. One of the microcentrifuge tubes was acidified (10 μL of 95% sulfuric acid) for the determination of ammonia (NH_3_) concentration using a colorimetric procedure based on the Berthelot (phenol-hypochlorite) reaction^55^. Absorbance was read at 550 nm using a Synergy2 plate reader (BioTek Instruments, Winooski, VT). The other microcentrifuge tube was not acidified and used for volatile fatty acid (VFA) concentrations, using a GC (7890A; Agilent Technologies Inc., Santa Clara, CA). This GC was equipped with a Nukol fused silica capillary column (30 m length × 0.25 mm I. D. × 0.25 µm d_f_, Supelco, Catalog #24107) for VFA separation according to procedures^56,57^. Briefly, samples were thawed at 4 °C and then centrifuged for 30-min at 700 × g. Then, 0.2 mL of 25% metaphosphoric acid containing 2-ethyl-butyrate as an internal standard was added, mixed, and chilled at 4 °C for 30-min. Samples were then centrifuged again at 5,000 × g for 30-min. A 0.8 mL subsample of the supernatant was aspirated, and 1.2 mL of methanol was added. A 0.45 µm polyethersulfone syringe filter was used to filter the sample and this was placed in a crimp top GC vial. The autosampler was programmed to inject 1 µL of sample into the split/splitless injector (at 250 °C) using a 50:1 split. The GC used helium (UHP grade) as a carrier gas set at a 1-mL/min flow rate. The GC oven had an initial temperature of 125 °C and was set to initially increase by 10 °C/min until it reached 175 °C followed by a 20 °C/min increase to 190 °C, for a total run time of 6.75 min.

All statistical analyses for the *in vitro* experiments were conducted using the R^58^ software (v4.4.1; R Core Team, 2024). Statistical significance was considered at *P* ≤ 0.10. All models included run (1-3) and gas production module as random effects. The treatment (control, imidazole, L-carnitine, methyl jasmonate, or propylpyrazine) were included as fixed effects. Gas production curves were fit to a non-linear model ^59^using the ‘*nlme*’ function^60^. Least square means were generated and means separation was conducted using the ‘*emmeans*’ package^61^. The same process was repeated for a second blind round (prior to metabolic modeling) for four more compounds dimethylglyoxime, propionyl-L-carnintine, pyrimidine, and safrole.

### 2.8 Microbial community metabolic modeling for understanding inhibitor-mediated flux shifts in rumen

Multi scale metabolic/ community model frameworks have proven effective in capturing inter organ and interspecies metabolic interactions and linking molecular level constraints to system level phenotypes^62,63^. We introduce an ensemble of Cowmunity1.0 metabolic models that reflect the steady-state ruminal flux distributions across control (no additives), and with-additives cases. Feed utilization in bovine rumen was predicted using the OptCom^64^ framework, a GAMS-based optimization platform for community metabolic modeling by *Zomorrodi et al*.^64^. To enable greater flexibility in model parsing and long-term code maintenance, the framework was translated into Python through the GAMSPy package. Draft genome-scale metabolic reconstruction was borrowed from *Islam et al*. (2019) and imported via XML parsing functions^65^, which converted the Systems Biology Markup Language (SBML)^66^ files into sets and variables compatible with GAMSPy. The reconstructions represented three key rumen microbial guilds: *Methanobrevibacter gottschalkii* (MGK, Archaea), *Prevotella ruminicola* (PRM, Bacteroidetes), and *Ruminococcus flavefaciens* (RFL, Firmicutes). In this preliminary demonstration, we have used equi-population mixtures of the three species. However, in our follow-up study we will incorporate 16S-RNA data to update the sub-population frequencies of each sub-species and add ancillary species such as protozoan ^67^(entodiniomorphs and holotrichs) and fungi^68^ (*Neocallimastigomycota*, *Basidiomycota*, and *Ascomycota*. Basidiomycota).

Constraints were incorporated to simulate metabolite uptake, and additional metabolites and reactions were introduced to better capture the known and predict new metabolic interactions among species. The model was optimized with the objective of maximizing total community biomass production. This optimized baseline model served as the control, against which treatment-simulated models were compared. Treatment simulations were generated by introducing new constraints derived from a literature review of relevant genes, binding affinity scores from docking to MCR, and metabolic reactions associated with the compounds of interest (details in supplementary information). While some molecules have known information on which pathways they impact in in the ruminal microbiome, this is not guaranteed for all. To this end, we provide a structo-metabolic strategy to identify any feed additive molecule, which microbial enzymes (across all species) are they likely to inhibit and consequently alter enzyme turnover kinetics and provide a structural theory of flux rewiring (reinforced using experimental measurements of volatiles). This will be stitched together in the follow-up, detailed metabolic dissection study.

### 2.9. Connecting the metabolism to structural biology

To elucidate the effect of treatment molecules on the metabolic pathways at the enzyme-metabolite level, we extracted a list of potential substrates to the enzymes present in the rumen microbial guild systems considered to conduct a similarity assessment of these substrates with the treatment molecules that putatively impair specific pathways that eventually result in methanogenesis flux changes. For each of the three systems metabolically modeled (MGK, PRM, and RFL), complete genome assemblies and all publicly available proteomes were retrieved from the NCBI taxonomy database^69^ (Taxonomy IDs: 1122229, 81582, and 1030842 respectively for MGK, PRM, and RFL). To obtain an overview of functional composition, the proteins were classified into broad functional categories based on sequence annotation keywords present in the proteome FASTA headers. These proteins descriptors were passed to identify annotations for enzymes, ion and metabolite transport proteins, nucleotide-binding proteins, regulatory proteins, and hypothetical/ unknown proteins, For the list of enzymes extracted from each species, we performed a search on the BRENDA^70^ database to retrieve EC numbers and associated ligands considering multiple matches including synonyms. These ligand names were mapped to corresponding Canonical SMILES via PubChem API ^71^ tool for similarity calculations. We use the size-torsion-chemistry aware ChemBERTa^43^ embeddings to calculate the similarity of the obtained ligands with each treatment molecule, allowing evaluation of potentially similar compounds that are associated with enzymes in downregulated pathways, as reported from the results of the metabolic model.

## 3. Results And Discussion

### 3.1. Structural analysis of MCR from Methanothermobacter marburgensis and known inhibitors

We employed the X-ray crystal structure of MCR (PDB ID: 5G0R) captured in its Ni(I)-methyl catalytic intermediate state, which represents a functional relevant snapshot of the enzyme during CH_4_ formation^72^. The methyl group of methyl-coenzyme M stated usually situates at least a 2.1 Å proximal to the Ni(I) of the MCR coenzyme F_430_ for a successful catalysis to materialize. There has been a proposal of reorganization of the substrate channel to bring the substrate species together; however, Ni (III)-methyl itself does not cause any noticeable structure change in the channel^2^. Given this, studies with biochemical and structural analysis of the MCR from *Methanothermobacter marburgensis* were focused on with the assumption that the last step of CH_4_ production in ruminants is the rate-limiting step of methanogenesis^73^.

A recent experimental study^2^ on the inhibitory properties of 3-NOP with the 3D structure of MCR (PDB ID: 5G0R) deciphered at a high resolution of 1.25 Å was selected for our study. In agreement with previous literature^29–31^, the MCR protein selected is a 273 kDa hexameric protein (**Figure 2**) with two catalytic grooves that are 50 Å apart. The MCR protein has a deep active site pocket with a substrate groove that runs ∼ 30 Å from the Ni(I) to the protein’s surface^1^. The activity of MCR, as reported by computational analysis from experimental data^72,73^, revealed the enzyme remains active only when its Ni ion is in the tetrapyrrole derivative of the cofactor F_430_ and has a +1-oxidation state, therefore catalyzing the last CH_4_-production step of methanogenesis in the rumen of livestock such as cattle, sheep, and goats^1,76,77^.

**Figure 2.**
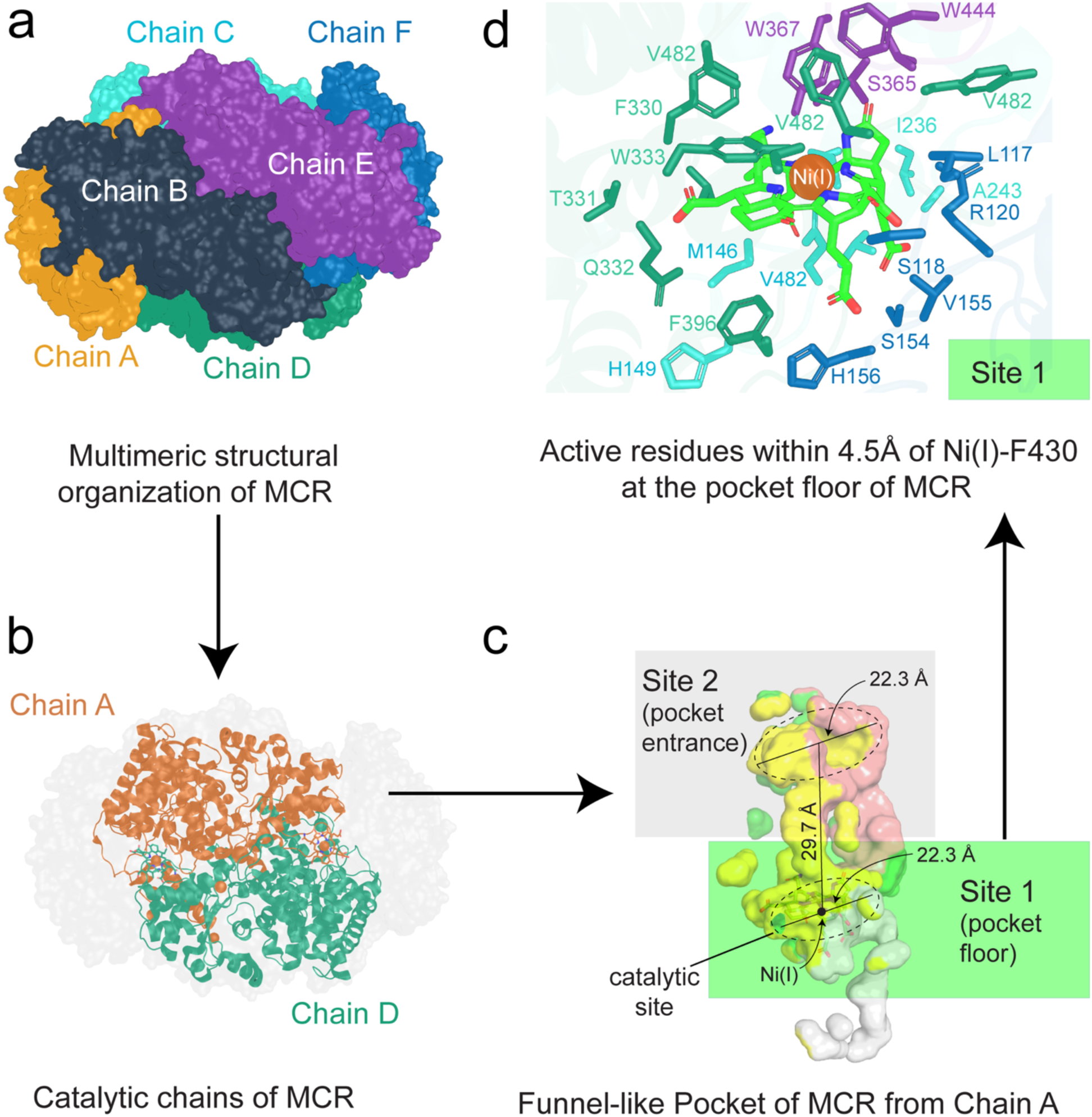
Illustration of the crystal structure of Methyl Coenzyme M Reductase (MCR) (PBD accession ID: 5G0R) from *Methanothermobacter marburgensis* and the six-chain hexameric complex. **a)** Each chain of MCR crystal structure has been indicated with six colors. **b)** Catalytically active chains (A and D) of MCR are shown in green and blue, while other non-catalytic chains are shown in gray. **c)** The funnel like pocket of MCR with F430 with site of binding designations. Site 1: 15Å from Ni(I) of F430 while Site 2: 16Å till surface of MCR from Ni(I) in F430. **d)** The active site residues of MCR within 4.5Å range of Ni(I) of F430.

All literature-based reported inhibitor compounds for enteric methanogenesis inhibition were collected. Distinct molecular compounds were selected, including statins, pterins, nitro-ol/esters, Coenzyme-B (CoBs), and CHBr_3_ and its analogs along with the native MCR substrate methyl-coenzyme M (CoM) (see **Figure 3**). Three statins (lovastatin, rosuvastatin, and simvastatin), four nitro-ol/esters (2-nitroethanol, 2-nitropropanol, 3-nitropropionate and 3-NOP), five coenzyme B analogs (CoBs) (N-5-mercaptopentanoylthreonine phosphate: CoB5, N-6-mercaptohexanoylthreonine phosphate: CoB6, N-7-mercaptoheptanoylthreonine phosphate: CoB7, N-8-mercaptooctanoylthreonine phosphate: CoB8, and N-9-mercaptononanoylthreonine phosphate: CoB9, and three pterins (pterin B53 (2,6-diamino-5-nitrosopyrimidin-4(3H)-one), pterin B54 (4-{3-[(2-amino-5-nitroso-6-oxo-1,6-dihydropyrimidin-4-yl)amino]propoxy}benzoic acid) and pterin B55 (2-amino-8-sulfanyl-1,9-dihydro-6H-purin-6-one) were studied along with CHBr_3_ and two other organobromines (bromopropionate and 2-bromoethanesulfonate [BES]) using detailed molecular modeling, thermodynamic assessment, and Ni(I) interactions in the presence of cofactor F_430_.

**Figure 3.**
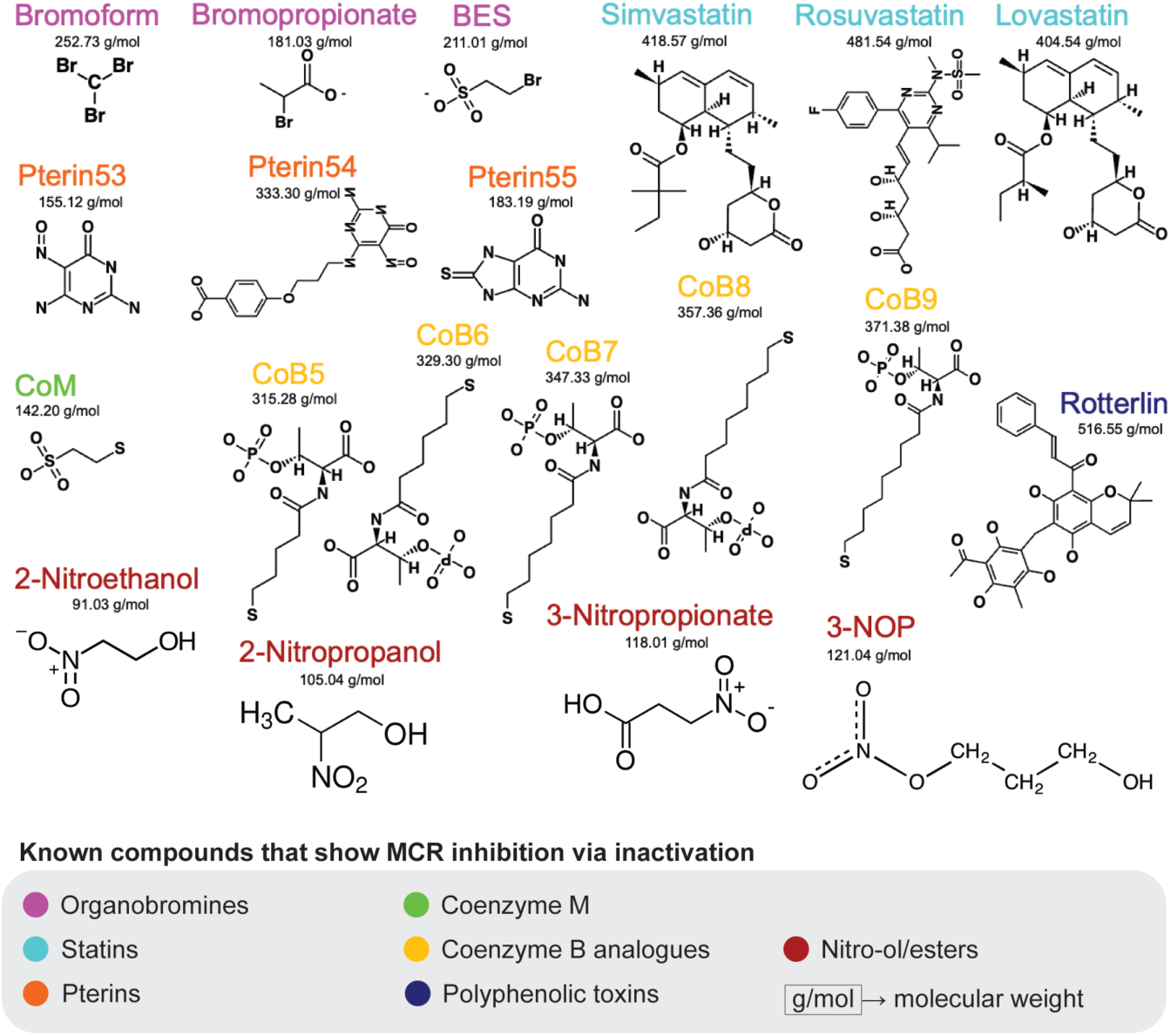
Structures of sixteen known anti-methanogenic molecules discussed in this study with their respective molecular weights. They are color-coded into groups organobromines (magenta): bromoform and its analogs, statins (blue), pterins (orange), coenzyme-M (green), coenzyme-B analogs (yellow), polyphenolic toxins (purple), and nitro-ol/esters (brown).

### 3.2 Pterins outperform other inhibitors in MCR binding affinity compared to bromoform

The top-scoring (strongest) binding poses were analyzed to evaluate the selected ligands’ binding affinities, interactions, and potential binding modes with no superimpositions within 5 Å. Conformational poses which have no van der Waal’s interaction with others were used to infer the maximum number of inhibitor molecules of each type that can simultaneously invade and yet remain biochemically bound within catalytic distances of Ni(I) of the cofactor F_430_ within the MCR enzyme pocket. The number of inhibitor molecules thus obtained is a representation of the maximum permissible stoichiometry of the inhibitor on a per-molecule basis of the MCR enzyme. Consequently, the inhibitor molecule poses that were accounted for were the ones within the electron transfer range with the Ni(I) of the tetrapyrrole of F_430_ in MCR.

Predicted binding affinities of known and candidate compounds to the Ni–F430 active site of MCR are summarized in **Figure 4** (part b). Across all molecules evaluated, docking scores spanned a wide energetic range of approximately –3 to –9 kcal·mol⁻¹, indicating an occurrence of weak, moderate, and high-affinity binders. Each value represents the mean of three independent docking replicates, demonstrating good intra-compound reproducibility.

**Figure 4:**
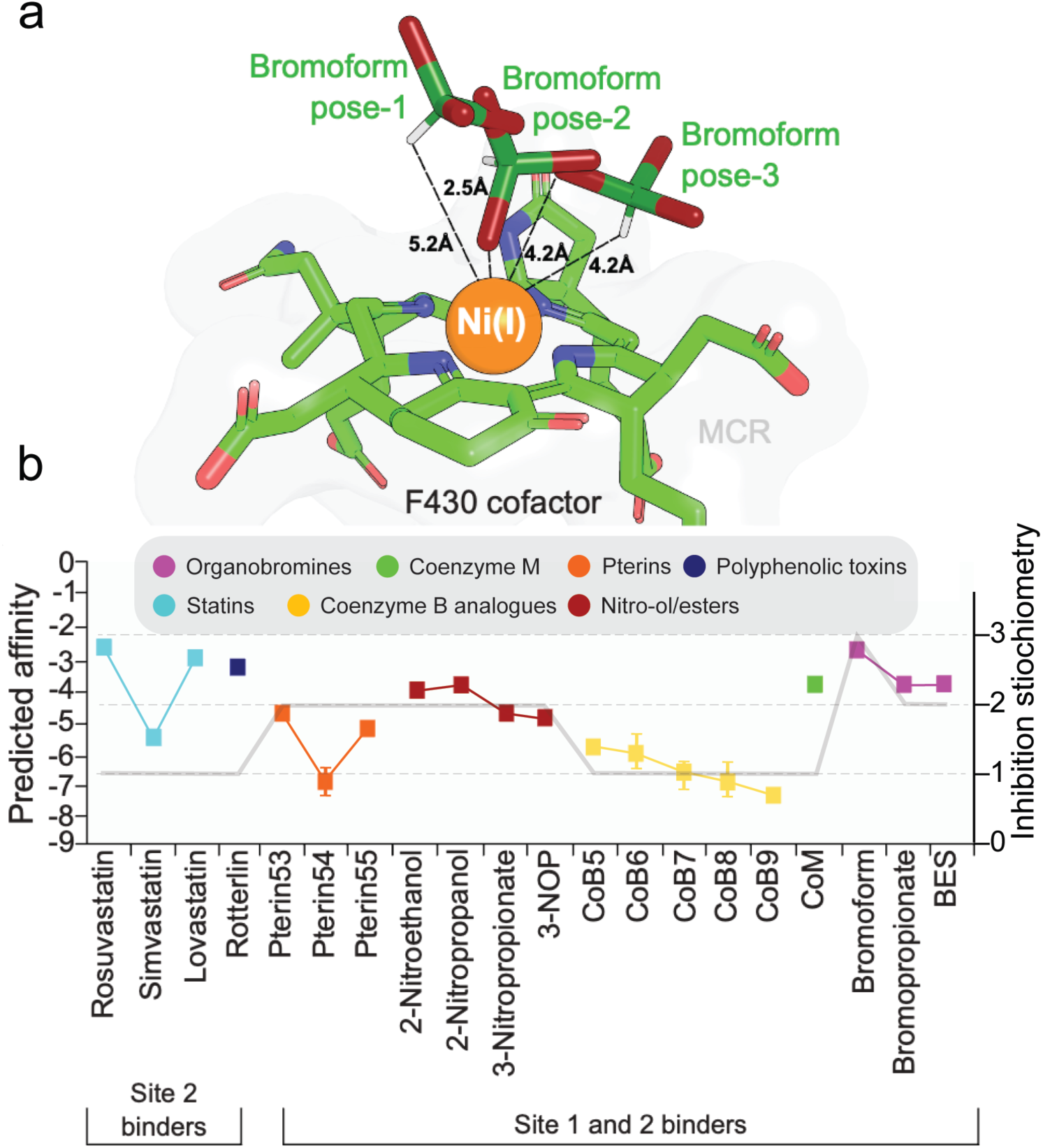
Atomic representation of the stoichiometric ratio of individual inhibitors docked to the active site of MCR enzyme in close vicinity of F_430_. **a)** Illustration of all three selected poses of bromoform interacting with Ni(I) of F_430_ in MCR protein. **b)** Scatter plot representation of the mean computationally predicted binding affinity values (only accounting for interactions with MCR enzyme and van der Waal’s with F_430_ and Ni(I)) of top three conformations of inhibitor molecules docked to F_430_ of MCR and also, a representation of all positive conformations of inhibitor molecules accurately posed within a 5 Å range.

Known anti-methanogenic compounds clustered in the moderate to strong affinity range (∼ –6 to –7.5 kcal·mol⁻¹). This group included nitro-alcohol derivatives such as 2-nitroethanol, 3-nitropropanol, and the commercially available inhibitor 3-NOP (trade name Bovaer, DSM-Firmenich, Maastricht, Netherlands), which are well-established reference inhibitors of methanogenesis. The correct energetic placement of these positive controls validates the predictive capacity of the docking workflow and confirms that the docking captures critical features of productive MCR–ligand interactions.

In addition, to other known inhibitors exhibited equal or greater predicted binding affinity than 3-NOP, with the strongest candidates approaching ∼ –9 kcal·mol⁻¹ such as CoB9 and Pterin54 (**Figure 4b**). These high-affinity ligands were predominantly drawn from the group classified as dual Site-1/Site-2 binders, suggesting that simultaneous engagement of multiple regions within the MCR catalytic pocket may contribute to inactivating MCR, hence CH_4_ inhibition. In contrast, molecules classified as Site-2-only binders tended to populate the weaker end of the affinity distribution.

Notably, the energetic separation between known inhibitors and their analogs provided a rational basis for MCR binding distribution amongst known inhibitors and their site of binding. Collectively, these results demonstrate that the energetics from molecular docking only enlighten the gray area of CH_4_ inhibition but sets foundational chemical diversity for deep learning-based prediction of safe metabolites in natural occurring databases and prior to *in vitro* fermentation screening.

### 3.3. MCR inhibitors cluster together when compared with ruminant specific metabolite databases

The quest to elucidate safe and available anti-methanogenic molecules from nature inspired the usage of the milk and bovine metabolome databases. However, the number of molecular features required to identify such anti-methanogenic molecules cluster is unknown due to the paucity of data in experimental literature. Thus, we chose to use a pre-trained SMILES embedding model called ChemBERTa, known primarily for molecular property predictions^43^.

The embeddings generated from the contrastive learning model were visualized using t-SNE plots (**Figure 5**) revealing clusters. The known anti-methanogenesis molecules and highly similar metabolites (positive class) formed compact clusters, while the remaining sample data (negative class) were more dispersed in embedding space. This highlighted the model’s efficiency in distinguishing the embedding features of anti-methanogenesis molecules from those of other metabolites.

**Figure 5.**
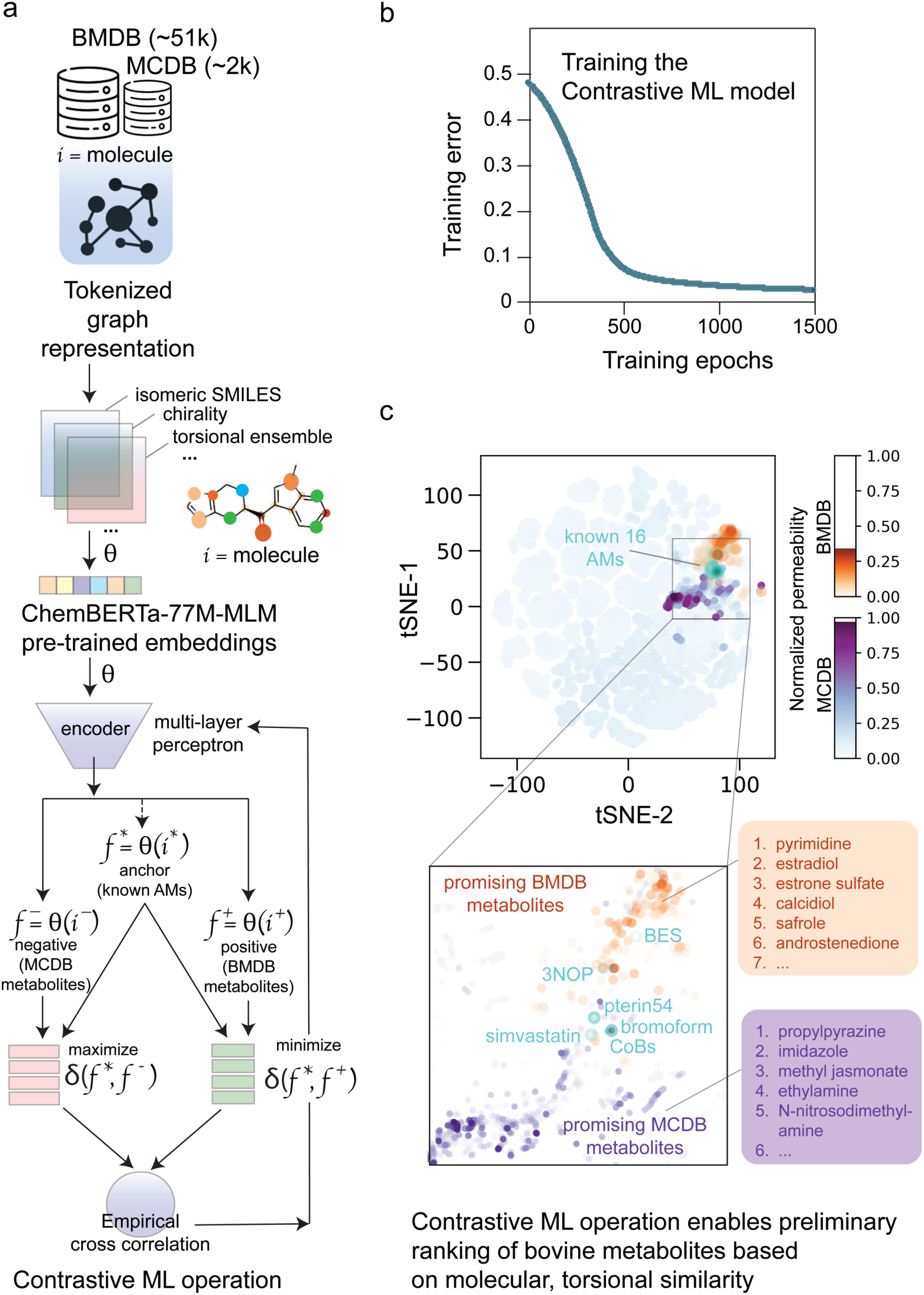
Two-dimensional t-SNE projection of molecular signatures reveals clustering of methanogenesis inhibitors. **a)** Workflow of contrastive learning for molecular feature from pre-trained model. **b)** Model training curve displaying training error per epoch. **c)** Visualization of clusters of embedding of known 16 anti-methanogenic molecules with the Milk Composition Database (MCDB) and Bovine Metabolome Database (BMDB). Color intensity signifies permeability of molecules (intense color means highly permeable). Numbered spots indicate list of some representative candidate molecules for further filtering.

To further enhance the analysis, the previously predicted permeability of each molecule was normalized (0-1) and incorporated as gradient colour intensities in the t-SNE visualization. The molecules with higher permeability are represented by darker intensities, enabling a visual correlation between permeability. The combination of permeability data with the t-SNE plots further enhanced interpretability, where one could spot molecules with structural and chemical likeness to known anti-methanogenesis compounds and yet possess favourable permeability features. This combined strategy effectively narrowed down the list of potential anti-methanogenesis molecules from the metabolite datasets, showcasing the utility of contrastive learning and visualization techniques in molecule discovery pipelines. Few representative molecules were numbered based on visualization (**Table 2**) as possible molecular targets.

**Table 2:** Representative molecules identified as a result from contrastive learning.

| Serial | MCDB | BMDB |
| --- | --- | --- |
| 1 | Pyrimidine | 3-hydroxyquinine |
| 2 | L-carnitine | 2,1-hydroxyprognenolone |
| 3 | Xestoaminol C | Aldosterone |
| 4 | Dimethylamine | Glutaryl carnitine |
| 5 | Methylamine | Propenoyl carnitine |
| 6 | (-)-alpha-pinene | 4-hydroxystachydrine |
| 7 | Androstenedione | 8-isoprostane |
| 8 | Estrone sulfate | Thromboxane |
| 9 | Fosfomycin | 14R,15S-EpETrE |
| 10 | Calcidiol | Cyanosulfurous acid anion |
| 11 | Estriol | N-Nitrosodimethylamine |
| 12 | Hexachlorocyclohexane | (S)-Methylmalonic acid semialdehyde |
| 13 |  | Dihydroceramide |

### 3.4. Investigation of predicted Anti-Methanogenic Molecules: Molecular Docking, Stoichiometric Perspectives, and MCR Inhibitory patterns

Using the MolToxPred model^50^, the toxicity scores were predicted after ranking and filtering down 1,111 clustered molecules using permeability values greater than 10 and Euclidean distances less than 10 to anti-methanogenic molecules candidates from the contrastive learning pipeline. **Table 3** highlights top 15 molecules from the filtered 1,111 molecules anti-methanogenic potential hence labelled predicted molecules (PMs). Molecular docking experiments for these top 15 molecules indicated that contrastive learning approach based on cosine similarity effectively captures potential anti-methanogenic candidates as an observed moderate binding from affinity scores was recorded. This finding strongly aligned with Yang *et al.* (2019) study demonstrating contrastive learning enhances molecular property prediction ^49,78^.

**Table 3:** Top 15 predicted anti-methanogenic molecules (PMs) based of Cellular Membrane Permeability and Euclidean distance from known anti-methanogenic molecules from literature.

| Predicted Molecules | Molecule Name | Affinity scores | Site of binding |
| --- | --- | --- | --- |
| PM1 | Isoquinoline | – 5.3 | 1 and 2 |
| PM2 | Safrole | – 6.7 | 1 and 2 |
| PM3 | Pyrimidine | – 3.9 | 1 and 2 |
| PM4 | Imidazole | – 3.0 | 1 and 2 |
| PM5 | Aniline | – 4.1 | 1 and 2 |
| PM6 | Nicotine-1'-N-oxide | – 4.1 | 1 and 2 |
| PM7 | Methyl jasmonate | – 7.8 | 1 and 2 |
| PM8 | (-)-Dihydrocarveol | – 4.9 | 1 and 2 |
| PM9 | Adrenosterone | – 3.6 | 1 and 2 |
| PM10 | Androstenedione | – 4.0 | 1 and 2 |
| PM11 | Aldrin | – 1.0 | 1 and 2 |
| PM12 | 6-Dimethylaminopurine | – 5.1 | 1 and 2 |
| PM13 | Propenoylcarnitine | – 4.2 | 1 and 2 |
| PM14 | L-Carnitine | – 4.0 | 1 and 2 |
| PM15 | Propylpyrazine | – 4.8 | 1 and 2 |

Out of 15 predicted molecules (PMs), the least binding affinity score recorded was −1 kcal·mol⁻¹, representing minimal interaction with the target (**Table 3**). On the contrary, some of the top-performing molecules exhibited significantly higher affinities, corroborating that binding free energy remains a key component in molecular recognition^79^ and MCR inactivation.

Stoichiometric analysis, thus quantifying flooding effect for inhibition, identified three non-clashing binding orientations of PM4 superimposing well with bromoform, the key known methanogenesis inhibitor. This similar conformational binding suggests chemical structure and spatial positioning of the inhibitors are vital to inhibit MCR enzyme, in agreement with previous research on CH_4_ inhibition mechanisms^80^. Further investigation questioned whether structural similarity was a predictor of MCR inhibition.

Our findings indicate that Euclidean distance is not a highly predictive factor of CH_4_ inhibition on a large scale as MCR inactivation by PMs poorly corelates in a linear fashion (r = 0.03*, p* = 0.911). Overall, this contradicts the prediction that structurally similar molecules possess similar biological activity^81^. Again, the fact that there is no high correlation reveals that molecular similarity alone is inadequate in predicting inhibitory potential. Instead, certain molecular interactions, hydrogen bonding patterns, and electrostatic complementarity may be more significant binding affinity determinants, as described in previous accounts in computational drug discovery research^82,83^. These results reveal the inadequacies of similarity-based screening methods, which in most instances rely on Euclidean distances or fingerprint-based clustering^84^. Our findings validate and as well supports the growing evidence that this deep learning-enabled method of ours (contrastive learning), can overcome such limitations by disconnecting non-linear molecular correlations^45,78,85^.

### 3.5. Prediction of Relative inhibitor binding probabilities and binding stability constants with Ni(I)

A scatter plot from the ChemBERTa similarity of compounds in BMDB, MCDB and the curated Sigma-Aldrich datasets to 3-NOP and their normalized permeability to 3-NOP revealed unique compounds of commendable permeability and similarity to 3-NOP (**Figure 6**).

**Figure 6.**
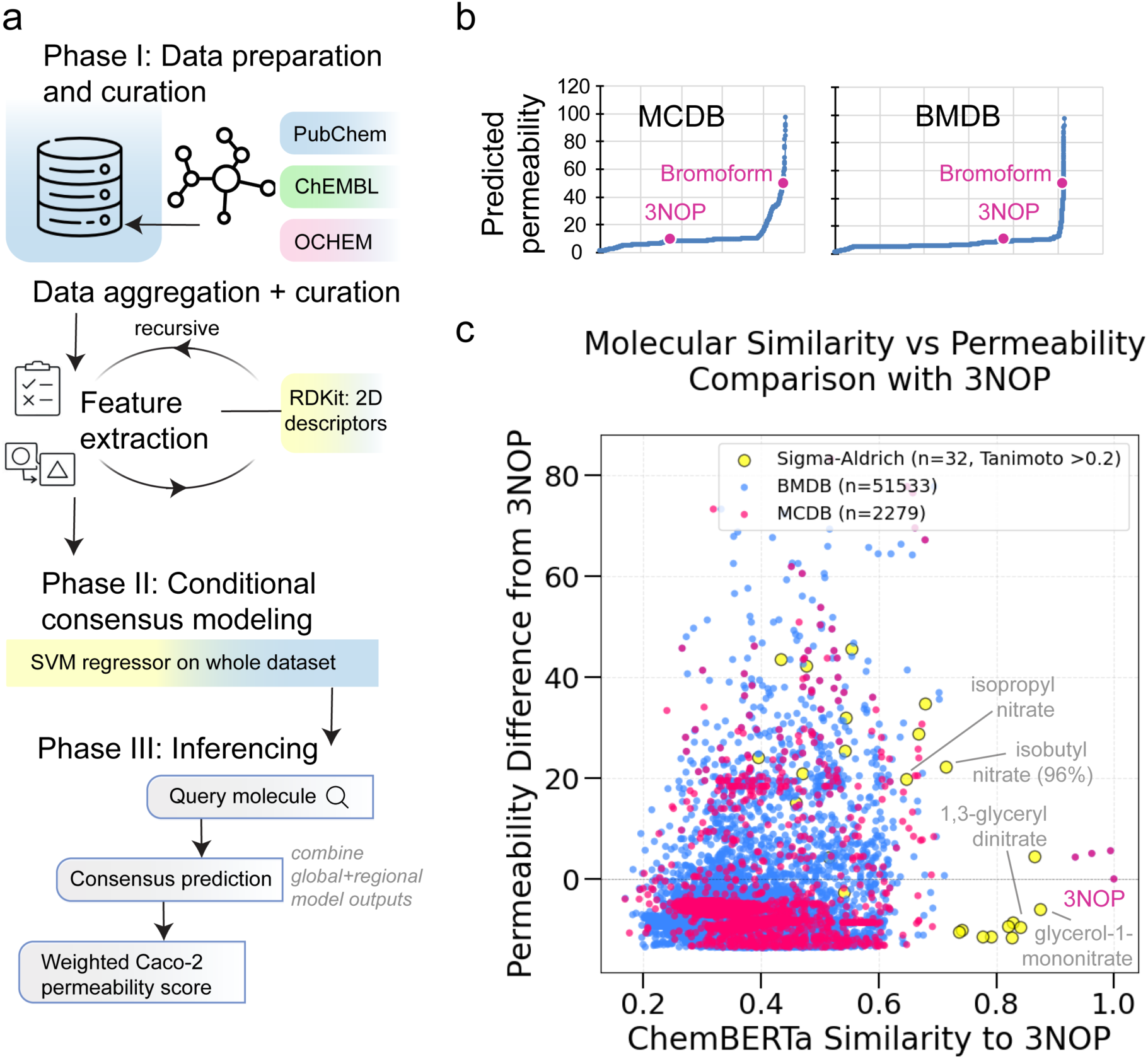
Structural similarity-permeability relationships constrain nitrate-derived analogs of 3-NOP. **a)** Flowchart representing the training and inference protocol for prediction of membrane permeability of small molecules. **b)** Predicted permeability of ∼2k MCDB metabolites, and ∼51k BMDB metabolites using the trained permeability predictor. All data points have been sorted in ascending order and then plotted; 3NOP and bromoform permeabilities are indicated. **c)** Scatter plot showing ChemBERTa structural similarity to 3-NOP (x-axis) versus relative membrane permeability difference compared with 3-NOP (y-axis) for compounds derived from Sigma-Aldrich (yellow, *n* = 36), BMDB (blue, *n* = 51,533), and the MCDB (red, *n* = 2,279).

These representative nitrate-containing molecules include isopropyl nitrate, isobutyl nitrate, glycerol-1-mononitrate, and especially 1,3-glyceryl dinitrate of the Sigma-Aldrich dataset highlights commercially available alternates to 3-NOP. Boltz-2 IC_50_ predictions for the above listed top-candidate inhibitor compounds to apo-MCR structure including 3-NOP (580.24 µM), Isopropyl nitrate (209.84 µM), Glycerol-1-mononitrate (163.57 µM), 1,3-Glyceryl dinitrate (72.10 µM), Isobutyl nitrate (16.17 µM), Pentaerythritol tetra nitrate (10.10 µM) support relative ranking of molecules, despite being agnostic to the presence of F_430_ cofactor chelating on Ni(I). Naive binding stability constant predictions for the above 5 compounds with Ni(I) on LOGKPREDICT resulted in nearly equal stability constants (fractional logK ranging from 0.99 to 1.01) to that of 3-NOP.

### 3.6. Bovine rumen fluid for in vitro testing of selected PMs validates computational uncertainty with methane inhibition

Four of the previously mentioned PMs were randomly chosen (as treatment molecules; TMs) for *in vitro* fermentation assays with basal substrate (**Table 1**) to ascertain CH_4_ reduction as a validation campaign for our established computational pipeline. Across all treatments, total gas production and CH_4_ output remained unchanged throughout the 48-h in vitro ruminal incubation. Twenty-four–h gas production ranged from 79.0 to 86.8 mL (*P* = 0.22), and 48-h gas production ranged from 100.3 to 109.0 mL (*P* = 0.29). Similarly, CH_4_ concentration was unaffected by treatment (3.79–4.10%; *P* = 0.84), as was CH_4_ production (3.85–4.52 mL; *P* = 0.33) as highlighted in **Table 4**. These results demonstrate that none of the evaluated compounds exerted direct inhibitory effects on enteric methanogenesis under the conditions tested.

**Table 4:** Effects of first round of 4 TMs on *in vitro* ruminal fermentation, CH_4_ production and VFA profiles.

| Item<sup>2</sup> | Treatment<sup>1</sup> |  |  |  |  |  |  |
| --- | --- | --- | --- | --- | --- | --- | --- |
| | Control | Imidazole | L-carnitine | Met J. | Prop Py. | SEm<sup>3</sup> | $P^4$ |
| initial pH | 6.653<sup>ab</sup> | 6.64<sup>b</sup> | 6.646<sup>b</sup> | 6.664<sup>a</sup> | 6.645<sup>b</sup> | 0.0266 | 0.06 |
| final pH | 6.295 | 6.299 | 6.295 | 6.300 | 6.300 | 0.0495 | 0.97 |
| IVTDMD, % | 70.36<sup>a</sup> | 69.26<sup>b</sup> | 69.86<sup>ab</sup> | 69.78<sup>ab</sup> | 69.66<sup>b</sup> | 0.884 | 0.05 |
| 24-h gas, mL | 79.01 | 86.36 | 82.55 | 84.28 | 86.84 | 4.744 | 0.22 |
| 48-h gas, mL | 100.30 | 108.18 | 104.98 | 106.399 | 109.00 | 4.456 | 0.29 |
| CH<sub>4</sub> conc., % | 3.79 | 3.83 | 3.88 | 3.89 | 4.10 | 0.644 | 0.84 |
| CH<sub>4</sub> prod., mL | 3.85 | 4.09 | 4.08 | 3.97 | 4.52 | 0.604 | 0.33 |
| Orskov param. |  |  |  |  |  |  |  |
| B, mL | 106.75<sup>c</sup> | 114.98<sup>a</sup> | 111.65<sup>b</sup> | 112.86<sup>b</sup> | 116.20<sup>a</sup> | 3.871 | <0.01 |
| C, %/h | 5.54<sup>b</sup> | 5.77<sup>a</sup> | 5.57<sup>ab</sup> | 5.68<sup>ab</sup> | 5.69<sup>ab</sup> | 0.064 | 0.08 |
| NH<sub>3</sub>, mmol/L | 26.33<sup>b</sup> | 29.1<sup>a</sup> | 28.96<sup>a</sup> | 28.80<sup>a</sup> | 29.49<sup>a</sup> | 3.96 | 0.06 |
| Acetate, mmol/L | 50.22 | 52.68 | 52.40 | 51.01 | 51.24 | 3.366 | 0.12 |
| Propionate, mmol/L | 26.74<sup>ab</sup> | 27.28<sup>a</sup> | 27.20<sup>a</sup> | 26.21<sup>b</sup> | 26.29<sup>b</sup> | 2.129 | 0.08 |
| Butyrate, mmol/L | 12.33 | 12.76 | 12.63 | 12.13 | 12.43 | 1.080 | 0.28 |
| Valerate, mmol/L | 2.99<sup>ab</sup> | 3.03<sup>a</sup> | 3.04<sup>a</sup> | 2.84<sup>c</sup> | 2.90<sup>bc</sup> | 0.664 | 0.07 |
| Isobutyrate, mmol/L | 1.09<sup>c</sup> | 1.18<sup>a</sup> | 1.15<sup>ab</sup> | 1.10<sup>bc</sup> | 1.12<sup>bc</sup> | 0.093 | 0.08 |
| Isovalerate, mmol/L | 2.95 | 3.13 | 3.08 | 2.94 | 2.98 | 0.329 | 0.13 |
| Total VFA, mmol/L | 96.32 | 100.06 | 99.52 | 96.23 | 96.98 | 6.135 | 0.14 |
| A:P, mmol/mmol | 1.89<sup>b</sup> | 1.95<sup>a</sup> | 1.95<sup>b</sup> | 1.96<sup>b</sup> | 1.96<sup>b</sup> | 0.140 | 0.01 |
| A+B:P, mmol/mmol | 2.36<sup>b</sup> | 2.42<sup>a</sup> | 2.42<sup>a</sup> | 2.42<sup>a</sup> | 2.43<sup>a</sup> | 0.190 | 0.07 |

<sup>1</sup> Met J. = methyl jasmonate; Prop Py. = propylpyrazine
IVTDMD = *in vitro* true dry matter digestibility; CH<sub>4</sub> conc = methane concentration; CH<sub>4</sub> prod = methane production; Orskov param = the nonlinear Orskov model parameters; B = asymptotic gas production; C = fractional rate of gas production; NH<sub>3</sub> = ammonia; VFA = volatile fatty acid; A:P = the acetate to propionate ratio; A+B:P = acetate plus butyrate to propionate ratio.
<sup>abc</sup> means with different superscripts differ ( $P \leq 0.10$ )
<sup>3</sup> SEm = standard error of the mean <sup>4</sup> The analysis of variance $P$ – value

In contrast, IVTDMD was modestly but significantly reduced (*P* = 0.05), declining from 70.36% in the control to 69.26% in the imidazole and propylpyrazine treatments, respectively. As gas production is tightly coupled to substrate degradability, the stability of total gas and CH_4_ output despite reduced IVTDMD indicates that CH_4_ was not suppressed through general microbial inhibition or loss of fermentative capacity. Instead, fermentative efficiency declined without a parallel reduction in methanogenic end products, a pattern inconsistent with effective CH_4_ inhibition.

Interestingly, we observed preserved microbial activity from fermentation kinetics. Nonlinear gas production modeling confirmed the absence of broad microbial suppression. The asymptotic gas pool (Orskov parameter B) differed significantly among treatments (*P* < 0.01), ranging from 106.8 to 116.2 mL, while the fractional gas production rate (C) showed an increase (*P* = 0.08). These shifts indicate that microbial fermentation capacity remained robust, ruling out microbial toxicity or generalized energetic suppression as it validates explanations for the lack of CH_4_ mitigation.

Uniquely, fermentation stoichiometry shifted toward hydrogen availability despite the unchanged concentration of total VFAs (96.2–100.1 mmol/L; *P* = 0.14) while acetate and butyrate remained stable, an indication that treatments effects emerged as primarily at the level of stoichiometric partitioning than total gas production. Again, the acetate-to-propionate (A:P) ratio increased significantly (*P* = 0.01) from 1.89 in the control to 1.95–1.96 across treatments. The combined acetate-plus-butyrate-to-propionate ratio (A+B:P) also increased (*P* = 0.07). Acetate and butyrate production generate molecular hydrogen (H₂), whereas propionate formation consumes H₂. Therefore, increases in A:P and A+B:P ratios reflect enhanced hydrogen availability to methanogenic archaea, providing a direct biochemical explanation for the absence of CH_4_ suppression^86^. Rather than redirecting reducing equivalents away from methanogenesis, these compounds promoted a fermentation architecture that favors hydrogen supply to CH_4_-forming pathways.

Isobutyrate concentration increased in treated groups (*P* = 0.08), rising from 1.09 mmol/L in the control to as high as 1.18 mmol/L in the imidazole treatment. Isobutyrate originates primarily from valine deamination, indicating enhanced amino acid fermentation^87,88^. This shift suggests greater ruminal nitrogen turnover and increased liberation of NH₃ and in fact, branched-chain VFA and NH_3_ concentrations are often positively correlated^89,90^. This supposition is confirmed by the increased (*P* = 0.06) NH_3_ concentrations by all treatments compared with the control.

The combined stability of CH_4_ output, preservation of fermentation kinetics, and elevation of hydrogen-producing fermentation pathways strongly indicate that none of the tested compounds inhibit MCR, the terminal CH_4_-forming enzyme in methanogenic archaea. Effective MCR inhibitors characteristically reduce CH_4_ independent of total gas production^91,92^ and often induce compensatory redirection of hydrogen toward propionate^92,93^ and hydrogen gas emission but neither of these mechanistic signatures was observed. Detailed flux information from round two of next four treatment molecules (dimethyl glyoxime, propionyl L-carnitine, pyrimidine, safrole) are listed in **Table S3.**

### 3.8. Microbial metabolic model outputs correlate with in vitro testing of predicted anti-methanogenic compounds

The Cowmunity1.0 model successfully reproduced baseline rumen community and provided mechanistic predictions for each treatment in methanogenesis (**Table 5**). Under controlled conditions, the optimized community exhibited a total biomass flux of 1.9376 gDCW·gDCW^-1^·h^-1^. Biomass contributions were distributed as follows: PRM dominated community growth by 1.4419 gDCW·gDCW^-1^·h^-1^, RFL contributed 0.2957 gDCW·gDCW^-1^·h^-1^, and MGK contributed 0.1999 gDCW·gDCW^-1^·h^-1^. Baseline CH_4_ production from MGK was 517.97 mmol·gDCW^-1^·h^-1^, consistent with model validation benchmarks. Introducing imidazole resulted in a slight decrease in total biomass with MGK biomass exhibiting a marginal increase (0.2010 gDCW·gDCW^-1^·h^-1^) but observed CH_4_ flux rose to 524.50 mmol·gDCW^-1^·h^-1^. In detail, imidazole treatment led to an increase in biomass production rates, substrate uptake, and metabolite output, consistent with its reported inhibition of lysosomal activity in predatory protozoa^94^. The model predicted a 0.36% increase in CH_4_ emissions, which did not fully capture the magnitude of the experimentally observed 1.05% increase. L- carnitine treatment decreased total biomass flux by 1.9093 gDCW·gDCW^-1^·h^-1^ and MGK biomass to 0.1779 gDCW·gDCW^-1^·h^-1^, reflecting reduced sterol export capacity. Despite this, CH_4_ production increased to 548.06 mmol·gDCW^-1^·h^-1^. Also, L-carnitine, was previously demonstrated to reduce cholesterol levels in cattle^95^, produced a simulated CH_4_ flux increase of 6.79%. This result was in closer agreement with the experimentally measured 2.37% increase. Also, methyl jasmonate and propylpyrazine increased CH_4_ flux to 552.63 mmol·gDCW^-1^·h^-1^ and 550.97 mmol·gDCW^-1^·h^-1^ respectively.

**Table 5:**
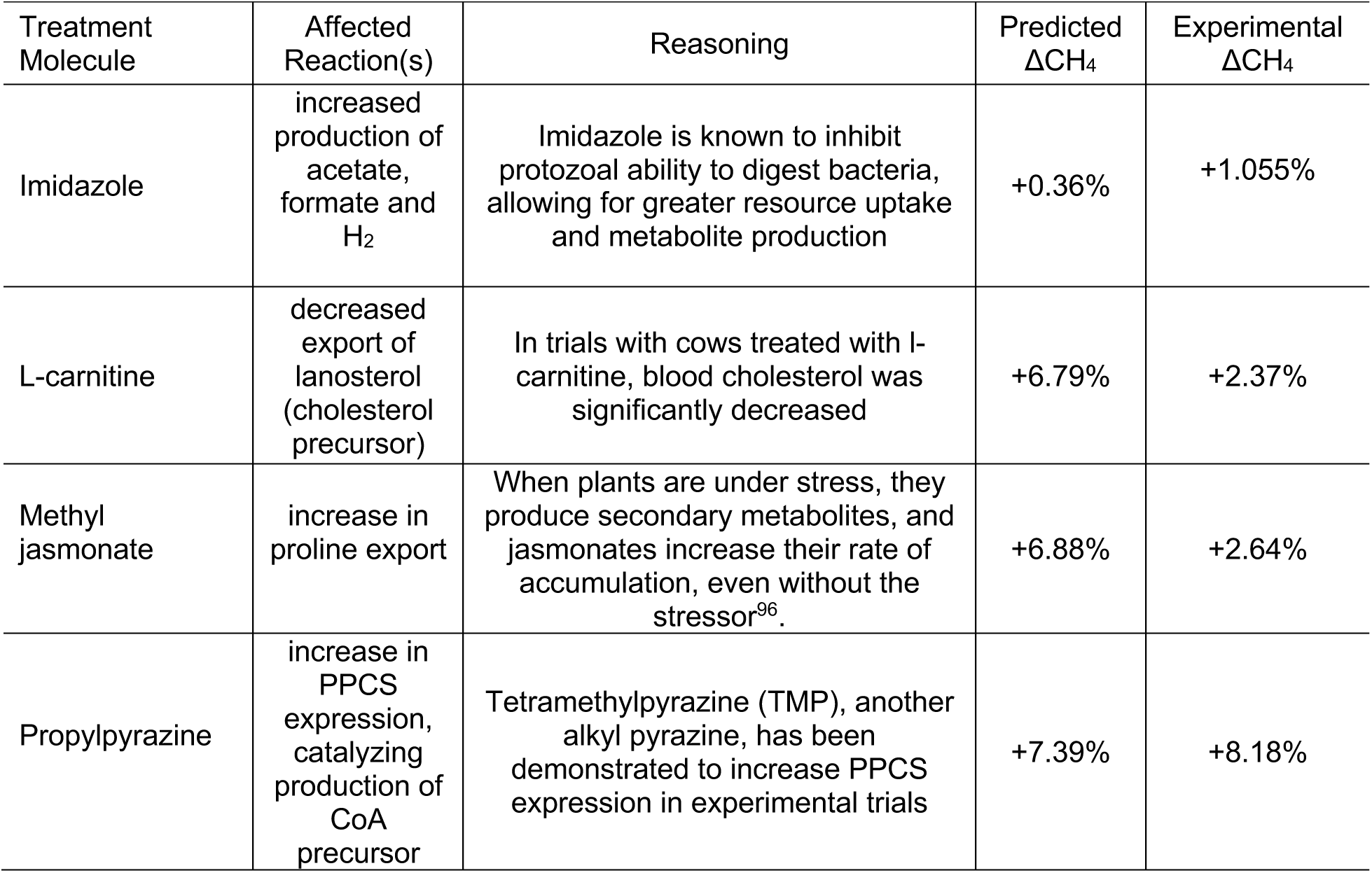
Predicted and experimental changes in CH_4_ emissions form our literature-based model.

| Treatment Molecule | Affected Reaction(s) | Reasoning | Predicted $\Delta\text{CH}_4$ | Experimental $\Delta\text{CH}_4$ |
| --- | --- | --- | --- | --- |
| Imidazole | increased production of acetate, formate and H<sub>2</sub> | Imidazole is known to inhibit protozoal ability to digest bacteria, allowing for greater resource uptake and metabolite production | +0.36% | +1.055% |
| L-carnitine | decreased export of lanosterol (cholesterol precursor) | In trials with cows treated with l-carnitine, blood cholesterol was significantly decreased | +6.79% | +2.37% |
| Methyl jasmonate | increase in proline export | When plants are under stress, they produce secondary metabolites, and jasmonates increase their rate of accumulation, even without the stressor<sup>96</sup>. | +6.88% | +2.64% |
| Propylpyrazine | increase in PPCS expression, catalyzing production of CoA precursor | Tetramethylpyrazine (TMP), another alkyl pyrazine, has been demonstrated to increase PPCS expression in experimental trials | +7.39% | +8.18% |

Here, methyl jasmonate, a phytohormone known to stimulate secondary metabolite production under stressed conditions, yielded a 6.88% computational increase in CH_4_ output (**Table 5**). This aligned moderately with the 2.64% increase observed from *in vitro* testing. Here in, propylpyrazine, an aromatic compound commonly found in foods such as coffee, demonstrated the greatest divergence between simulation and experiment. While its direct biological effects are not well established, literature on related alkylpyrazines particularly tetramethylpyrazine (TMP) shows that TMP can enhance expression of the *ppc* gene which has been associated with an artificial serine cycle to enable the bioconversion of both CH_4_ and CO_2_ into succinate^97^ resulted in approximately 17% of CH_4_ increase^98^. In the present study, propylpyrazine treatment resulted in a 7.39% increase in CH_4_ emissions computationally, compared to a much larger 8.18% increase measured experimentally.

### 3.9. Structo-metabolic analyses of small molecule feed additives

A similarity analysis of the putative inhibitor ligands to the enzymes in the studied metabolic guild (MGK, PRM, and RFL) with the treatment molecules indicated a wide distribution from 20% to 100% on the chemical + sequence similarity space indicated by the cosine similarity of ChemBERTa embeddings (**Figure 7**). We extracted a list of metabolites associated with downregulated reactions in the three metabolic models that had more than 70% cosine similarity and mass difference less than 250 Da relative to the treatment molecules and mapped them to 95 unique enzymes, predominantly involved in energy metabolism, amino acid metabolism and glycolysis pathways.

**Figure 7.**
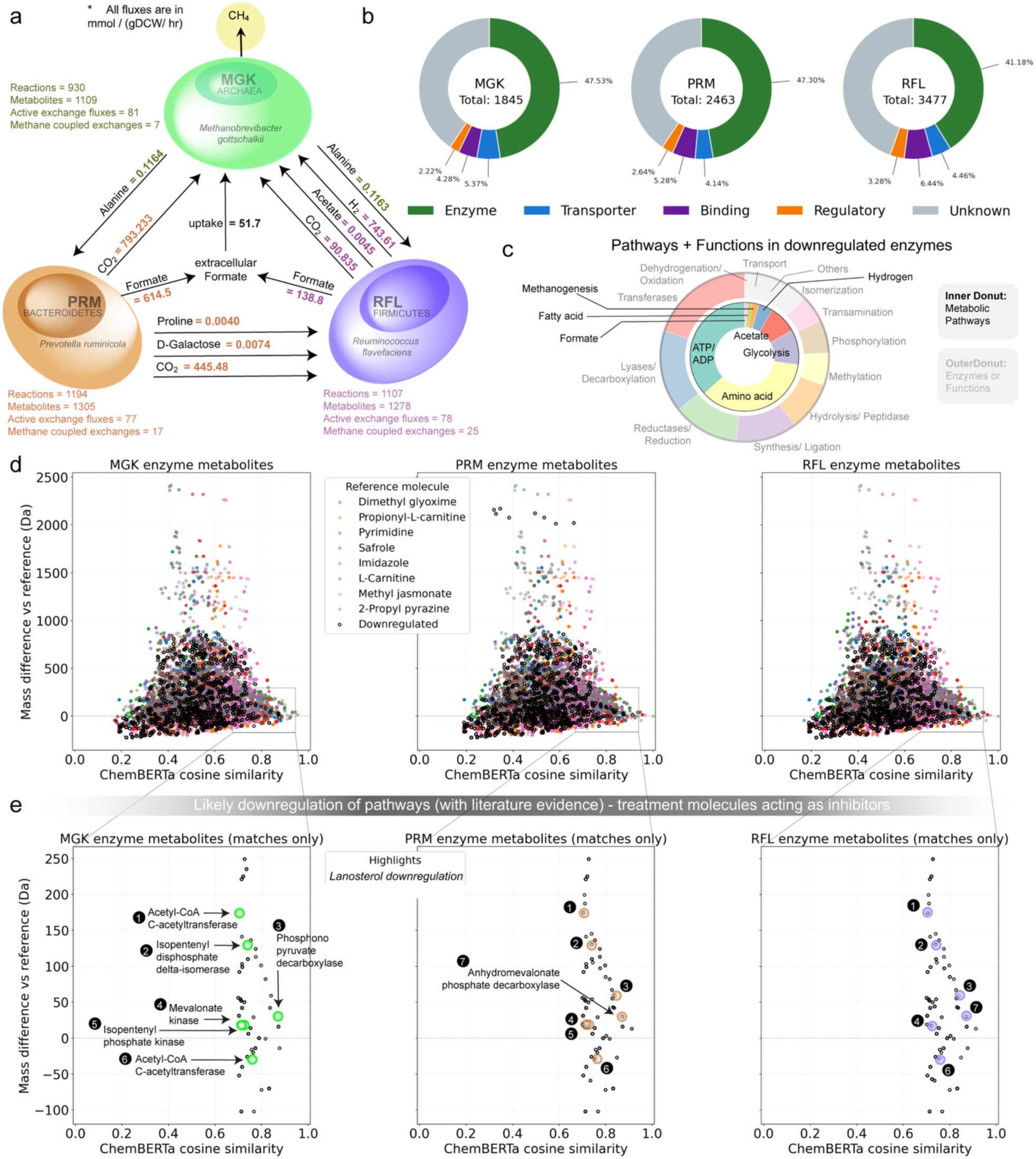
**a)** Schematic representation of the three member Cowmunity metabolic model showing major interspecies metabolite exchanges among *Prevotella ruminicola* (PRM), *Ruminococcus flavefaciens* (RFL), and *Methanobrevibacter gottschalkii* (MGK). **b)** Functional composition of the proteome space in rumen metabolic species used in the Cowmunity model. **c)** Pathways and functions associated with 95 unique enzymes in downregulated metabolites with more than 70% ChemBERTa cosine similarity and 250 Da mass difference to the treatment molecules **d)** Scatter plot showing ChemBERTa cosine structural similarity versus relative mass difference with 8 treatment molecules (dimethyl glyoxime, propionyl-L-carnitine, pyrimidine, safrole, imidazole, L-carnitine, methyl jasmonate, and 2-propyl pyrazine) for enzyme metabolites involved in metabolic pathways in all three species. **e)** Potential inhibitor metabolites with >75% ChemBERTa cosine similarity containing 7 critical metabolites involved in downregulated lanosterol biosynthesis.

A set of all genomes present in the three rumen metabolic organisms considered in our metabolic guild (namely MGK, PRM and RFL) available on the NCBI taxonomy database allowed identification and classification of the corresponding proteome space into proteins that potentially act as enzymes, ion and metabolite transporters, regulatory proteins, and nucleotide binding proteins. Since the eight treatment molecules considered for the Cowmunity model (dimethyl glyoxime, propionyl L-carnitine, pyrimidine, safrole, imidazole, L-carnitine, methyl jasmonate and 2-propyl pyrazine) are now known to exhibit high bacterial membrane permeability, the overall flux distribution of reactions and metabolite levels targeted for reduced methanogenesis would ultimately depend on the inhibitory role of associated metabolites in the enzymes, which point to upregulated/downregulated pathways.

We obtained a list of plausible ligand substrates to all enzymes present in the three systems by mapping them to all available native metabolite substrates on BRENDA known to interact with enzymes in the same family as that of the query enzyme in the metabolic model. The metabolites in downregulated pathways, as predicted by our Cowmunity1.0 model, that match with the list of ligand substrates analyzed for similarity to the treatment molecules. Such enzyme metabolites with a high chemical and size similarity to the treatment molecules indicate potential inhibition tendencies. **Figure 8** shows a wide distribution of size and chemistry aware ChemBERTa embedding cosine similarities of the enzyme metabolites of varying molecular masses with the treatment molecules. We extracted those metabolites common to all three species with more than 70% ChemBERTa cosine similarities and within 250 Da mass difference from any of the 8 treatment compounds considered. We identified enzymes associated with these metabolites and mapped them to their metabolic pathways, including enzymes metabolite pairs involved in the lanosterol biosynthesis pathway (as an exemplar) **Figure 8**, which is downregulated in feed-restricted cows^99^ .

For these high similarity inhibitors compounds and the associated enzymes, the relative inhibition efficiency can be estimated using RealKcat^100^ to widen the scope of search towards obtaining a recipe for functionally positive methanogenesis inhibition.

This analysis is useful even in cases where there are no literature priors on the upregulation/downregulation of pathways to study the effect of treatment molecules on the flux distribution and overall methanogenesis inhibition performance. Specifically, by learning pathway-specific metabolite-similarity thresholds we can build estimates on which treatment molecules are likely to inhibit which pathways preferentially. For example, L-carnitine administration *in vitro* resulted in downregulation of sterol biosynthesis pathway (experimentally reported to lower lanosterol levels by ∼11%) primarily in MGK and RFL species. This is corroborated from existing literature that L-carnitine treatment demonstrate potential to drop cholesterol levels^101,102^. **Figure 7** shows that as many as seven enzymes in related pathways spanning all three microbial species can be inhibited (partially) by L-carnitine owing to its >70% similarity to their native substrates. This threshold can be leveraged for new treatment molecules to predict expected sterol downregulation ability and consequently affect methane levels. Subsequently, permeability estimations combined with molecular dynamics trajectories for microbial transporters of chosen inhibitor metabolites will allow ranking of treatment molecules for effective movement across the cytosolic membrane for subsequent shortlisting.

## 4. Conclusion

MCR inhibition has the greatest potential to mitigate enteric CH_4_ emission from ruminant livestock. Here, we computationally compared known inhibitor molecules reported to be explored as MCR inhibitors. The key takeaway of this work is not to demonstrate that a single round of experiments with eight treatment molecules is sufficient to discover a singular “*Rosetta Stone*” molecule that outperforms commercial 3NOP, nor does it argue that eight treatments constitute an endpoint of search. Instead, we establish a rigorously interpretable, structurally and metabolically grounded workflow that converts limited—often mislabeled as “negative”—experimental data into mechanistic insight. At a time when machine-learning-driven biology prioritizes best-case, often inconsequential state-of-the-art metrics, we interrogate eight non-optimal molecules as informative probes, revealing digestion enhancement, methane-flux modulation, and MCR–F430-Ni(I) interaction signatures for the first time, that together define a reusable, scalable, multi-scale design framework for methane mitigation.

Through molecular docking, we showed that some known inhibitor molecules have relatively higher binding affinity to MCR than the well-known CHBr_3_ and nitro-ol/ester compounds, which are the key CH_4_ inhibitory molecules of choice. We also revealed that the reaction dynamics and the overall mechanistic understanding of the inhibition process is greatly influenced by the stoichiometry of the inhibitors in the active site as demonstrated by the flooding effect of CHBr_3_. Specifically, the presence of three bromine atoms in bromoform makes it a highly effective halogenated compound for competitively inhibiting the interaction of natural substrates with the Ni(I) ion in the F_430_ cofactor in MCR enzyme. Notably, inhibitor stoichiometry does not only dictate the binding affinity as a factor for CH_4_ inhibition but also the extent of methyl transfer inhibition and, consequently, the reduction in CH₄ release. We demonstrate that the stoichiometry of the inhibitors in the active site, as deduced from the non-superimposing docking poses within the active site groove, is directly proportional to the size of the inhibitor. It can be interpreted that smaller inhibitors have higher flooding effects within the active site. Also, the contrastive learning inspired t-SNE clustering indicated that all the known 16 inhibitor molecules explored in this study have inherent similarities among themselves when compared to ruminant specific metabolites and reveal some potential candidates from these databases as anti-methanogenic agents and their precursors. However, *in vitro* rumen incubations of four selected molecules failed to suppress CH₄ production despite modest effects on fermentation stoichiometry. Consistent with these findings, community metabolic modeling revealed that redirection of hydrogen and VFA partitioning can counteract putative direct MCR inhibition. Collectively, these results demonstrate that molecular similarity, docking affinity, and permeability predictions must be integrated with community- and fermentation-level biology to reliably predict CH_4_ mitigation. This work establishes a scalable discovery framework while underscoring the necessity of multi-scale validation prior to field deployment. Next in future iterations, we will (i) adopt an active-learning strategy in which experimentally tested TMs serve as labeled positives and hard negatives to recalibrate a multi-task model that predicts methane suppression, feed efficiency, and nitrogen losses; (ii) augment molecular representations with mechanistically informed descriptors derived from MCR-F_430_ structural dynamics; and (iii) deploy reinforcement learning and contextual bandit formulations to prioritize, identify and possibly generate compounds that maximize a composite reward integrating these phenotypes. In this way, the present negative findings become an asset, shaping a more biologically grounded discovery engine for enteric methane-mitigation agents.

## Associated Content

### Supporting Information

The Supporting information is compiled and available free of charge at the link to be added later.

### Code availability

All code used for molecular docking, contrastive learning and community metabolic modeling is publicly available at GitHub: https://github.com/ChowdhuryRatul/burpblockers.git. The repository includes scripts, figures and analyses reported in this study. Also, other files used for chemical space investigation in Figure 7 can be found here: https://huggingface.co/datasets/chowdhury-lab/3nop-similarity-excel-files/tree/main.

### Author Contributions

The project was conceived by RC. The simulations and analyses were set up and performed by RA. AB and MSN, together developed and performed the contrastive learning pipeline. ZRM performed cowmunity metabolic modeling and analysis for this study. KS and SN helped in data collection and assisted with simulations. VSR performed binding affinity prediction of apo-MCR and inhibitor compounds. SD conducted Tanimoto similarity analysis and Caco-2 permeability studies. TJM provided a valuable discussion on the competitive inhibition of enzymes, which guided the study. MB, NF, and JK provided valuable feedback which helped in designing the study and supervised conducting the *in vitro* gas production experiments. RA and RC wrote the manuscript. All authors helped in editing the manuscript. No authors declare any competing interests. All authors agree with this final version of the manuscript.

## ACKNOWLEDGMENT

R.C. acknowledges support through Iowa State University Startup Grant, Building A World of Difference Faculty fellowship, CIRAS Applied Mini-Grant, and NSF 22-599, EPSCoR RII Track-1, Award Number DQDBM7FGJPC5 for partially funding this study. R.A. was funded by NIH R35GM143074 to T.J.M. T.J.M. is also supported by the Karen and Denny Vaughn Faculty Fellowship. J.A.K. was supported by the USDA ARS Project # 3090-31630-006-000D (Strategies to Manage Feed Nutrients, Reduce Gas Emissions, and Promote Soil Health for Beef and Dairy Cattle Production Systems of the Southern Great Plains). Mention of company or trade names is for description only and does not imply endorsement by the USDA. USDA is an equal opportunity provider and employer. The authors will also love to appreciate Lana Castleberry and Bridgette Hiltbrunner (both at USDA-ARS) for assisting in conduction in vitro gas production experiments as well as Evan Ingmire and Swathi Nadendla for their extensive literature search on candidate methanogens with MCR activity. We also acknowledge the use of Google Scholar for the significant literature search process as it greatly contributed to collation of relevant references used in this study.

## Animal and Human Care and Use Approval

Procedures for collecting ruminal fluid from cannulated steers were approved by the West Texas A&M University Institutional Care and Use Committee (#2022.01.001).

## ABBREVIATIONS

3-NOP: 3-nitrooxypropanol
AFOLU: Agriculture, Forestry and Other Land Use
BMDB: Bovine Metabolome Database
CHBr_3_: bromoform
CH_3_-S-CoM: methyl-coenzyme
M CO_2_-e: CO_2_-equivalence
CoB5: N-5-mercaptopentanoylthreonine phosphate
CoB6: N-6-mercaptohexanoylthreonine phosphate
CoB7: N-7-mercaptoheptanoylthreonine phosphate
CoB8: N-8-mercaptooctanoylthreonine phosphate
CoB9: N-9-mercaptononanoylthreonine phosphate
CoB-HS: methyl-coenzyme B
DM: dry matter
D-MPNN: directed message passing neural network
F_430_: coenzyme F_430_
GHG: greenhouse gas
IPCC: Intergovernmental Panel on Climate Change
IVTDMD: *in vitro* true DM digestibility
ipTM: interface predicted TM-score
MCDB: Milk Composition Database
MCR: Methyl Coenzyme M Reductase
PM: predicted molecule
PRM: *Prevotella ruminicola*
RCSB PDB: Research Collaboratory for Structural Bioinformatics Protein Data Bank
ReLU: rectified linear unit
SBML: Systems Biology Markup Language
SDF: Structure-Data File
SMILES: Simplified Molecular Input Line Entry System
TM: Template Modeling

**Nonstandard Abbreviations**
A:P: Acetate-to-propionate ratio
Adam optimizer: adaptive Moment Estimation.

