## Supporting Information for "Structural and Metabolic Characterization of Ni(I)-inhibitors Provide a Robust Anti-Methanogenicity Scoring System"

**KEYWORDS:** enteric methanogenesis, bromoform, enzyme inhibition, emissions mitigation, climate change, contrastive learning, Cowmunity modeling.

***Active site description of Methyl coenzyme M reductase***

Methyl coenzyme M reductase (MCR) catalyzes the reduction of methyl-coenzyme M (CoM-HS) in the gut of bovine rumen to produce methane ($CH_4$) as a natural means of reducing excess hydrogens for energy production. This reaction only proceeds in the presence of a key oxidized state of Ni, a component of cofactor $F_{430}$ situated in the active site groove of MCR enzyme. The MCR enzyme has a hexameric ($\alpha_2\beta_2\gamma_2$) chained crystal structure with two symmertric catalytic sites. One active site situated in chain A while the other in chain D. As such, the catalytic site studied in this study was based on chain A yet the groove which is made up of chain A, D and C were jointly studied. Key active site residues that form the active pocked were determined by estimation of all residues within 4.5Å of coenzyme $F_{430}$ of MCR (PDB ID: **5G0R)** in PyMOL software.

***Molecular Docking***

Selected inhibitor molecules collected from literature were docked using AutoDock Vina[1,2]. Following the rigid docking procedure, MCR and its native coenzyme $F_{430}$ were kept rigid while inhibitor molecules were made to freely move in the system. Grid box coordinates x, y and z were the coordinates of Ni(I) ion in the structure with size of 30x30x30 to focus on the cofactor $F_{430}$ of MCR. Binding poses from the results were screened with a non-superimposing criterion as well as poses within 4.5 Å distance from the Ni(I) of $F_{430.}$ Below are the top selected poses of inhibitor molecules within 4.5 Å proximal range of Ni(I) of cofactor F30 within the active site groove of MCR used in this study with their affinity values compared to COM (see **Table S1**).

**Table S1**: Energy scores of top 3 binding poses and their mean for all inhibitors.

| Type of Molecule | Computational Binding Energy scores | | | |
|---|---|---|---|---|
| | Pose A | Pose B | Pose C | Mean |
| Rosuvastatin | -2.78 | -2.78 | -2.78 | -2.78 |
| Simvastatin | -5.76 | -5.76 | -5.76 | -5.76 |
| Lovastatin | -3.91 | -3.91 | -3.91 | -3.91 |
| Rottlerin | -3.31 | -3.31 | -3.31 | -3.31 |
| Pterin53 | -4.94 | -4.86 | -4.84 | -4.88 |
| Pterin54 | -7.726 | -7.66 | -6.16 | -7.17 |
| Pterin55 | -5.30 | -5.26 | -5.30 | -5.28 |
| 2-Nitroethanol | -4.30 | -4.22 | -3.87 | -4.13 |
| 2-Nitropropanol | -4.27 | -3.92 | -3.77 | -3.99 |
| 3-Nitropropionate | -5.37 | -4.84 | -4.62 | -4.94 |
| 3-NOP | -5.39 | -5.28 | --4.53 | -5.07 |
| Bromoform | -2.88 | -2.86 | -2.50 | -2.75 |
| Bromopropionate | -4.06 | -3.99 | -3.71 | -3.92 |
| BES | -4.42 | -3.80 | -3.51 | -3.91 |
| COB5 | -6.42 | -6.03 | -5.78 | -6.08 |
| COB6 | -6.72 | -6.58 | -5.25 | -6.18 |
| COB7 | -7.32 | -7.01 | -6.14 | 6.83 |
| COB8 | -7.33 | -7.67 | -6.33 | -7.11 |
| COB9 | -7.07 | -7.67 | -7.51 | -7.63 |
| COM* | -4.45 | -3.80 | -3.59 | -3.95 |

Molecular docking affinity scores of all known inhibitors normalized by that of native MCR substrate (CoM) affinity score reveal the only simvastatin, all pterins, 3-nitropropionate, 3-NOP and coenzyme B analogs have substantial affinity to MCR-$F_{430}$-Ni(I) (**Figure S1**).

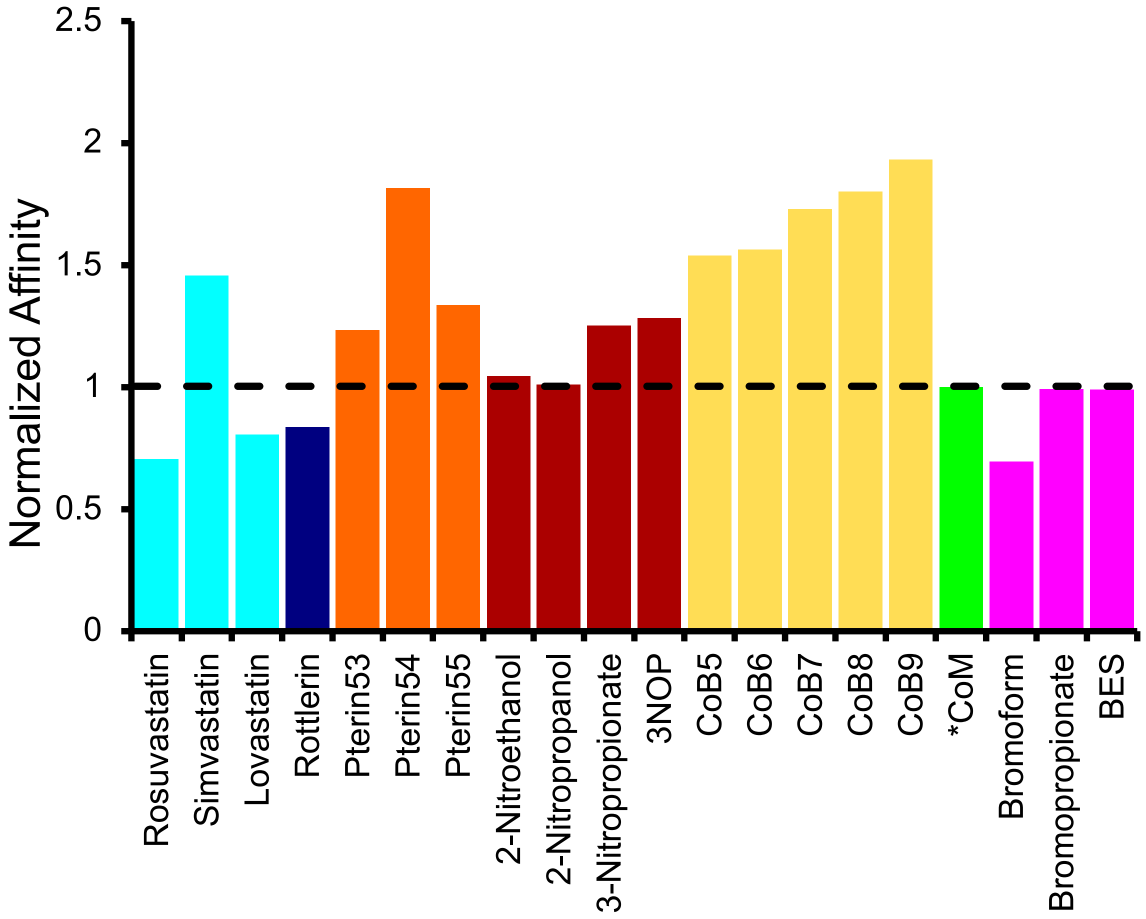

**Figure S1**: Normalized binging affinity scores of known inhibitor molecules to CoM. Molecules are colored in groups: Statins (cyan), Phenolic compound (dark blue), Pterins (orange), Nitro-ol/esters (maroon), Coenzyme B analogs (yellow), Methyl-coenzyme M (green) and Organobromines (pink).

***Cowmunity metabolic modelling detailed methods***

A three species genome scale Cowmunity model was developed to simulate metabolic interactions within bovine rumen. The model integrates *Methanobrevibacter* (MGK), *Prevotella rumincola* (PRM), and *Ruminococcus flavefaciens* (RFL) as species for MCMR inhibition (**Figure S2**). PRM and RFL degrade plant polysaccharides (cellulose, D-glucose, D-galacturonate) and release key fermentation products ($H_2$, $CO_2$, formate, acetate, alanine). MGK consumes $H_2$, $CO_2$, and formate to drive methanogenesis, producing $CH_4$ as the terminal output.

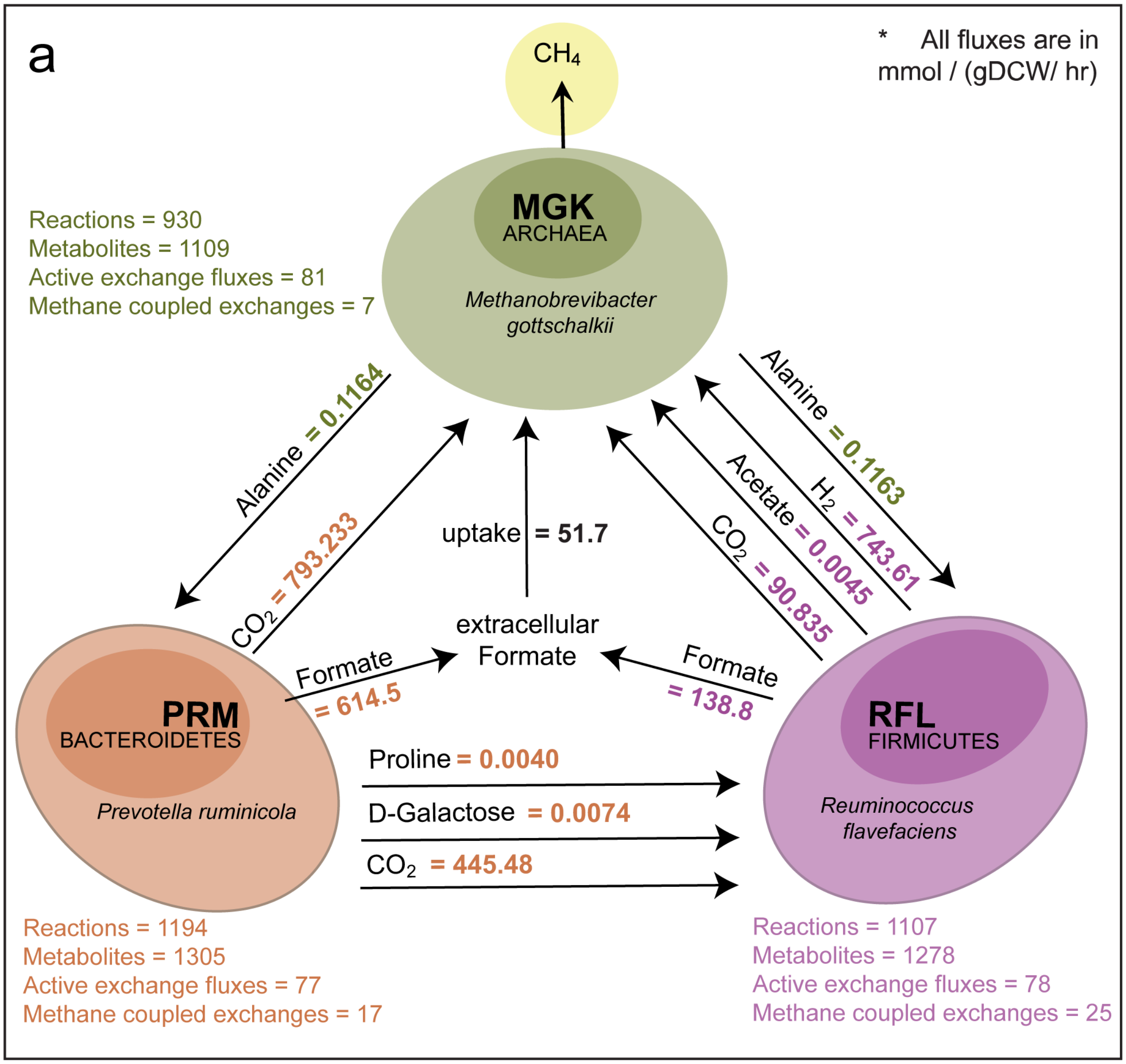

**Figure S2:** Schematic representation of the three member Cowmunity metabolic model showing major interspecies metabolite exchanges among *Prevotella ruminicola* (PRM), *Ruminococcus flavefaciens* (RFL), and *Methanobrevibacter gottschalkii* (MGK).

The Species SBML models were acquired from prior study[3], and curated to standardize exchange reactions and remove duplicate metabolites. All SBML files were parsed using libSBML[4], and corresponding reaction or metabolite sets were generated in GAMSPy[5] to create the computational container. Each organism was assigned a stoichiometric matrix (S), reaction set (J), and metabolite set (I). Metabolic fluxes ($V$) were constrained by reaction reversibility and maximum bounds (±1,000 mmol·gDCW$^{-1}$·h$^{-1}$). The steady state mass balance was imposed for every organism as

$$S.V = 0$$

Community interactions were represented using explicit transfer variables that quantify the movement of extracellular metabolites between species. For each shared metabolite $m((H_2, CO_2, formate, acetate, sugars, amino\ acids)$ we defined:

$EX_m^k$: the exchange flux of metabolite $m$ for species $k$, where positive values indicate secretion and negative values indicate uptake. $trans_m^k$: the inter species transfer variable denoting the amount of metabolite $m$ made available for consumption by species $k$ indicating the three community members: MGK, PRM and RFL. Transfer feasibility was enforced by requiring that the amount of metabolite available to MGK could not exceed the total secretion from the other two bacterial species:

$$trans_m^{MGK} \leq EX_m^{PRM} + EX_m^{RFL}$$

To ensure consistency between transfer and uptake, MGK's transfer variable was directly coupled to its own exchange flux:

$$trans_m^{MGK} = EX_m^{MGK}$$

Similar constraints were applied for PRM and RFL. The multi species optimization followed by OptCom[6] bilevel structure, consisting of an inner problem of maximizing each organism's biomass and also, an outer problem of maximizing the total community biomass:

$$\max biomass_{outer} = v_{MGK}^{biomass} + v_{PRM}^{biomass} + v_{RFL}^{biomass}$$

All constraints were implemented in GAMSPy, and the final model was solved as a nonlinear programming problem by IPOPT[7].

To incorporate treatments in the Cowmunity metabolic model, molecular docking against methyl coenzyme M reductase (MCMR) was performed using AutoDock Vina[1,2]. Docking inputs used for this model included four treatments (imidazole, L-cartinine, methyl jasmonate, and propylpyrazine) tested during in vitro gas production studies with bovine rumen fluid. Molecular docking simulation affinity scores (kcal·mol$^{-1}$) were converted to the dissociation constant $K_d$. The $K_d$ constant was converted into a quantitative fractional enzyme occupancy defined as:

$$f = \frac{[drug]}{[drug] + K_d}$$

Drug concentration in the rumen was assumed to be $0.01 \mu M$, because treatment compounds administered in feed undergo substantial dilution in the rumen fluid volume (about 100-150 L). Based on typical feed additive dosing rates and expected solubility limits, resulting in rumen concentrations generally fall within low micromolar range.

To transfer binding affinity from docking into a usable form for flux constraint modification, we defined a base inhibition term $I_{base}$. This term provides a standardized numerical representation of binding strength across different orders of magnitude of $K_d$. Rather than modeling kinetic inhibition explicitly, this term functions as a scaled indicator of how strongly a ligand is predicted to interfere with the MCR catalytic site. Strong binders are assigned values close to1, whereas weak binders receive values closer to 0.1. The resulting value serves as the foundational component used later in the workflow to compute the overall inhibition strength applied to reaction flux bounds.

$$I_{base} = \begin{cases} 0.995 & K_d < 0.001 \\ 0.98 + (log_{10}K_d + 3)(0.95 - 0.98), & 0.001 \leq K_d \leq 0.01 \\ 0.90 + \frac{0.1 - K_d}{0.09} 0.05, & 0.01 \leq K_d \leq 0.1 \\ 0.70 + \frac{1.0 - K_d}{0.9} 0.20, & 10^{-1} \leq K_d \leq 1 \\ 0.40 + \frac{10 - K_d}{9} 0.30, & 1 \leq K_d < 10^1 \\ 0.10 + \frac{100 - K_d}{90} 0.30, & 10 \leq K_d < 10^2 \\ 0.10, & K_d \geq 10^2 \end{cases}$$

$$I_{strenght} = f.I_{base}$$

A global biological relaxation factor was applied:

$$I_{relaxed} = I_{strength}.R_{global}$$

With $R_{global} = 0.15$. Also, a specific treatment relaxation was included:

$$R_{effective} = \frac{R_{global}}{R_{molecule}}$$

The final inhibition factor applied to flux bounds was:

$$IF = \max(0.005, 1 - I_{Strength}.R_{effective}.C$$

For competitive inhibitions, we scaled upper bounds only while non-competitive inhibitions had both upper and lower bounds scaled up. Reasoning behind this was competitive inhibition reduces the forward catalytic capacity but does not affect intrinsic reversibility hence only the upper bound needed to be scaled. On the other hand, non-competitive inhibition reduces the enzyme's overall activity, independent of direction, requiring both upper and lower flux limits to be reduced.

***Cowmunity metabolic modelling results***

The community model successfully reproduced baseline rumen community and provided mechanistic predictions for each treatment in methanogenesis (**Table S2**). Under control conditions, the optimized community exhibited a total biomass flux of 1.93 mmol·gDCW$^{-1}$·h$^{-1}$. Biomass contributions were distributed as follows: PRM dominated community growth by 1.4419 gDCW·gDCW$^{-1}$·h$^{-1}$, RFL contributed 0.2957 gDCW·gDCW$^{-1}$·h$^{-1}$, and MGK contributed 0.1999 gDCW·gDCW$^{-1}$·h$^{-1}$. Baseline methane production from MGK was 517.97 mmol·gDCW$^{-1}$·h$^{-1}$, consistent with model validation benchmarks. Introducing imidazole resulted in a slight decrease in total biomass (1.9361 gDCW·gDCW$^{-1}$·h$^{-1}$), but MGK biomass increased marginally (0.2010 gDCW·gDCW$^{-1}$·h$^{-1}$), also methane flux rose to 524.50 mmol·gDCW$^{-1}$·h$^{-1}$. L-carnitine treatment decreased total biomass to 1.9093 gDCW·gDCW$^{-1}$·h$^{-1}$ and MGK biomass to 0.1779 gDCW·gDCW$^{-1}$·h$^{-1}$, reflecting reduced sterol export capacity. Despite this, methane production increased to 548.06 mmol·gDCW$^{-1}$·h$^{-1}$. Also, methyl jasmonate and propylpyrazine increased methane flux to 552.63 mmol·gDCW$^{-1}$·h$^{-1}$and 550.97 mmol·gDCW$^{-1}$·h$^{-1}$ respectively.

**Table S2:** Predicted methane production and biomass fluxes in the Cowmunity Model under control and treatment conditions.

| Treatment | Methane Flux (mmol·gDCW$^{-1}$·h$^{-1}$) | Total Biomass Flux (gDCW·gDCW$^{-1}$·h$^{-1}$) | MGK Biomass Flux (gDCW·gDCW$^{-1}$·h$^{-1}$) |
|---|---|---|---|
| Control | 517.9731 | 1.937558 | 0.199912 |
| Imidazole | 524.499046 | 1.936136 | 0.200988 |
| L-Carnitine | 548.062346 | 1.909250 | 0.177922 |
| Methyl Jasmonate | 552.631884 | 1.932386 | 0.197516 |
| Propylpyrazine | 550.973123 | 1.937558 | 0.199912 |

Methane emission flux corresponds to the MGK-catalyzed methyl-coenzyme M reductase reaction. Total biomass flux represents community-level growth, while MGK biomass flux reflects methanogen-specific growth. All treatments produced higher methane flux than the untreated control, consistent with experimentally observed trends.

***Mapping metabolism to proteome space***

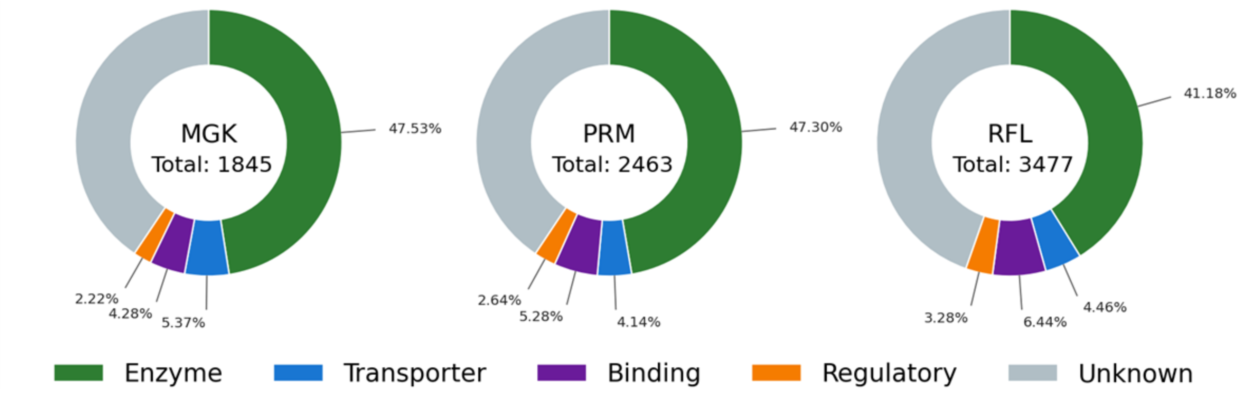

**Figure S3:** Functional composition of the proteome space in rumen metabolic species considered in this study.

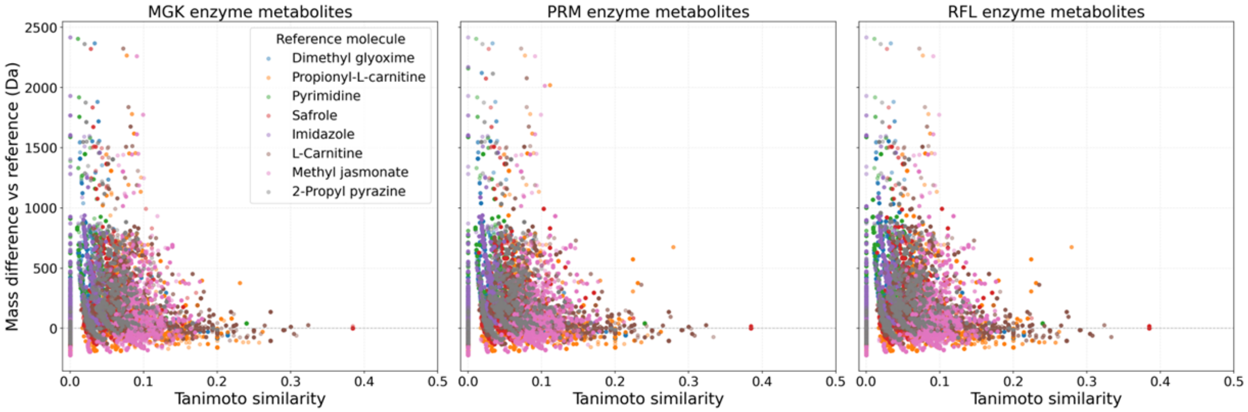

**Figure S4:** Tanimoto similarity versus mass difference of enzyme metabolites in the rumen species relative to the treatment molecules.

**Table S3:** Detailed flux information from round two of next four treatment molecules.

| Run | Bottle | Module | TRT | F57 BagIDs | initial.pH | final.pH | app.DMD |
|---|---|---|---|---|---|---|---|
| 1 | 14 | 32 | Control | 29_30 | 6.690 | 6.263 | 51.085 |
| 1 | 24 | 31 | Control | 11_12 | 6.680 | 6.265 | 51.303 |
| 1 | 9 | 47 | Control | 13_14 | 6.692 | 6.278 | 50.018 |
| 1 | 19 | 40 | Control | 15_16 | 6.674 | 6.275 | 49.275 |
| 1 | 13 | 49 | Dimethylglyoxime | 27_28 | 6.690 | 6.263 | 51.247 |
| 1 | 10 | 36 | Dimethylglyoxime | 5_6 | 6.687 | 6.263 | 49.050 |
| 1 | 17 | 38 | Dimethylglyoxime | 39_40 | 6.695 | 6.249 | 52.230 |
| 1 | 22 | 37 | Dimethylglyoxime | 17_18 | 6.702 | 6.272 | 50.923 |
| 1 | 7 | 29 | Propionyl-L-Carnitine HCl | 7_8 | 6.711 | 6.247 | 51.284 |
| 1 | 12 | 44 | Propionyl-L-Carnitine HCl | 1_2 | 6.680 | 6.274 | 49.165 |
| 1 | 1 | 25 | Propionyl-L-Carnitine HCl | 9_10 | 6.686 | 6.270 | 50.7630218 |
| 1 | 11 | 41 | Propionyl-L-Carnitine HCl | 31_32 | 6.708 | 6.287 | 50.5228843 |
| 1 | 16 | 34 | Pyrimidine | 21_22 | 6.689 | 6.371 | 49.8681333 |
| 1 | 3 | 39 | Pyrimidine | 23_24 | 6.682 | 6.268 | 49.7382228 |
| 1 | 21 | 45 | Pyrimidine | 3_4 | 6.682 | 6.274 | 50.7871784 |
| 1 | 6 | 48 | Pyrimidine | 25_26 | 6.703 | 6.267 | 49.4237957 |
| 1 | 15 | 33 | Safrole | 37_38 | 6.685 | 6.258 | 50.8522054 |
| 1 | 20 | 35 | Safrole | 19_20 | 6.683 | 6.249 | 50.3634873 |
| 1 | 4 | 43 | Safrole | 33_34 | 6.692 | 6.286 | 49.2343221 |
| 1 | 5 | 26 | Safrole | 35_36 | 6.682 | 6.256 | 50.8017723 |
| 2 | 4 | 39 | Control | 29_30 | 6.520 | 6.156 | 51.401222 |

| | | | | | | | |
|---|---|---|---|---|---|---|---|
| 2 | 1 | 41 | Control | 3_4 | 6.554 | 6.166 | 51.101887 |
| 2 | 10 | 46 | Control | 9_10 | 6.516 | 6.135 | 53.2679391 |
| 2 | 18 | 40 | Control | 1_2 | 6.532 | 6.128 | 50.6970508 |
| 2 | 17 | 37 | Dimethylglyoxime | 31_32 | 6.550 | 6.126 | 50.6474645 |
| 2 | 8 | 26 | Dimethylglyoxime | 39_40 | 6.534 | 6.152 | 48.9231466 |
| 2 | 13 | 35 | Dimethylglyoxime | 25_26 | 6.532 | 6.144 | 50.4601495 |
| 2 | 19 | 32 | Dimethylglyoxime | 5_6 | 6.540 | 6.160 | 50.5573432 |
| 2 | 20 | 27 | Propionyl-L-Carnitine HCl | 33_34 | 6.533 | 6.143 | 51.9392583 |
| 2 | 3 | 47 | Propionyl-L-Carnitine HCl | 7_8 | 6.568 | 6.135 | 51.6399093 |
| 2 | 24 | 38 | Propionyl-L-Carnitine HCl | 17_18 | 6.528 | 6.105 | 53.1695588 |
| 2 | 6 | 29 | Propionyl-L-Carnitine HCl | 35_36 | 6.520 | 6.133 | 51.8300772 |
| 2 | 16 | 33 | Pyrimidine | 11_12 | 6.513 | 6.140 | 50.6883824 |
| 2 | 12 | 36 | Pyrimidine | 37_38 | 6.555 | 6.147 | 50.909883 |
| 2 | 15 | 43 | Pyrimidine | 15_16 | | | |
| 2 | 21 | 25 | Pyrimidine | 21_22 | 6.537 | 6.125 | 52.4089926 |
| 2 | 14 | 48 | Safrole | 13_14 | 6.534 | 6.143 | 52.3659204 |
| 2 | 7 | 49 | Safrole | 19_20 | 6.534 | 6.189 | 51.0694542 |
| 2 | 5 | 34 | Safrole | 23_24 | 6.545 | 6.116 | 51.2838754 |
| 2 | 23 | 45 | Safrole | 27_28 | 6.566 | 6.152 | 49.6779319 |
| 3 | 19 | 29 | Control | 33_34 | 6.593 | 6.275 | 51.0133635 |
| 3 | 23 | 47 | Control | 39_40 | 6.638 | 6.253 | 51.1064703 |
| 3 | 18 | 25 | Control | 35_36 | | | |
| 3 | 13 | 38 | Control | 17_18 | 6.690 | 6.262 | 60.0054248 |
| 3 | 10 | 28 | Dimethylglyoxime | 5_6 | 6.669 | 6.254 | 53.8677763 |
| 3 | 16 | 48 | Dimethylglyoxime | 31_32 | 6.621 | 6.273 | 52.0901368 |
| 3 | 11 | 45 | Dimethylglyoxime | 29_30 | 6.674 | 6.270 | 52.9613937 |
| 3 | 24 | 49 | Dimethylglyoxime | 15_16 | 6.609 | 6.269 | 51.3962682 |
| 3 | 4 | 27 | Propionyl-L-Carnitine HCl | 13_14 | 6.607 | 6.273 | 51.9329868 |
| 3 | 15 | 37 | Propionyl-L-Carnitine HCl | 3_4 | 6.690 | 6.224 | 55.8144168 |
| 3 | 21 | 34 | Propionyl-L-Carnitine HCl | 37_38 | 6.667 | 6.257 | 54.3046143 |
| 3 | 2 | 31 | Propionyl-L-Carnitine HCl | 21_22 | 6.680 | 6.272 | 52.7251786 |
| 3 | 20 | 39 | Pyrimidine | 23_24 | 6.664 | 6.273 | 50.9436689 |
| 3 | 3 | 22 | Pyrimidine | 25_26 | 6.666 | 6.257 | 52.4325366 |
| 3 | 5 | 32 | Pyrimidine | 19_20 | 6.598 | 6.264 | 52.112969 |
| 3 | 22 | 44 | Pyrimidine | 11_12 | 6.689 | 6.265 | 53.8756267 |
| 3 | 6 | 41 | Safrole | 9_10 | 6.688 | 6.278 | 52.3592501 |
| 3 | 8 | 36 | Safrole | 27_28 | 6.671 | 6.273 | 52.5463496 |
| 3 | 14 | 35 | Safrole | 1_2 | 6.680 | 6.244 | 54.1850994 |
| 3 | 9 | 26 | Safrole | 7_8 | 6.585 | 6.275 | 51.6589525 |

| Run | Bottle | Module | TRT | F57 BagIDs | IVTDMD | CH4.per | 24h.gas.prod |
|---|---|---|---|---|---|---|---|
| 1 | 14 | 32 | Control | 29_30 | 69.095 | 4.9637 | 83.8826249 |
| 1 | 24 | 31 | Control | 11_12 | 68.370 | 5.0625 | 90.2219326 |
| 1 | 9 | 47 | Control | 13_14 | 67.747 | 5.21683333 | 84.7882403 |
| 1 | 19 | 40 | Control | 15_16 | 67.238 | 4.95983333 | 89.3163172 |
| 1 | 13 | 49 | Dimethylglyoxime | 27_28 | 68.917 | 5.19603333 | 83.3166153 |
| 1 | 10 | 36 | Dimethylglyoxime | 5_6 | 66.863 | 5.0602 | 82.0713941 |
| 1 | 17 | 38 | Dimethylglyoxime | 39_40 | 69.816 | 5.3016 | 88.7503076 |
| 1 | 22 | 37 | Dimethylglyoxime | 17_18 | 68.726 | 5.25716667 | 88.2974999 |
| 1 | 7 | 29 | Propionyl-L-Carnitine HCl | 7_8 | 69.708 | 5.27863333 | 86.1466634 |
| 1 | 12 | 44 | Propionyl-L-Carnitine HCl | 1_2 | 68.830 | 4.8389 | 89.7691249 |
| 1 | 1 | 25 | Propionyl-L-Carnitine HCl | 9_10 | 68.7185244 | 4.73773333 | 90.1087307 |
| 1 | 11 | 41 | Propionyl-L-Carnitine HCl | 31_32 | 66.8953973 | 5.16776667 | 78.4489326 |
| 1 | 16 | 34 | Pyrimidine | 21_22 | 67.6239418 | 5.08926667 | 81.9581922 |
| 1 | 3 | 39 | Pyrimidine | 23_24 | 68.8072281 | 4.80666667 | 88.4107018 |
| 1 | 21 | 45 | Pyrimidine | 3_4 | 67.2513638 | 5.19083333 | 82.0713941 |
| 1 | 6 | 48 | Pyrimidine | 25_26 | 68.4993703 | 4.9853 | 86.5994711 |
| 1 | 15 | 33 | Safrole | 37_38 | 68.4767636 | 4.9917 | 83.8826249 |
| 1 | 20 | 35 | Safrole | 19_20 | 67.9104041 | 4.68896667 | 88.4107018 |
| 1 | 4 | 43 | Safrole | 33_34 | 67.2967129 | 4.92116667 | 82.9770095 |
| 1 | 5 | 26 | Safrole | 35_36 | 68.775584 | 5.07283333 | 83.769423 |
| 2 | 4 | 39 | Control | 29_30 | 66.976373 | 4.68633333 | 72.6756345 |
| 2 | 1 | 41 | Control | 3_4 | 66.6788835 | 5.1246 | 72.1096249 |
| 2 | 10 | 46 | Control | 9_10 | 67.8876738 | 5.03246667 | 79.0149422 |
| 2 | 18 | 40 | Control | 1_2 | 67.5787491 | 5.14416667 | 79.9205576 |
| 2 | 17 | 37 | Dimethylglyoxime | 31_32 | 66.3331794 | 4.82193333 | 73.5812499 |
| 2 | 8 | 26 | Dimethylglyoxime | 39_40 | 66.0281863 | 4.68016667 | 72.2228268 |
| 2 | 13 | 35 | Dimethylglyoxime | 25_26 | 66.7184978 | 4.98786667 | 69.5059807 |
| 2 | 19 | 32 | Dimethylglyoxime | 5_6 | 66.2363684 | 4.78073333 | 73.5812499 |
| 2 | 20 | 27 | Propionyl-L-Carnitine HCl | 33_34 | 67.5078012 | 5.20856667 | 75.8452884 |
| 2 | 3 | 47 | Propionyl-L-Carnitine HCl | 7_8 | 67.0624507 | 4.53083333 | 73.5812499 |
| 2 | 24 | 38 | Propionyl-L-Carnitine HCl | 17_18 | 68.4475107 | 5.08455 | 85.3542499 |
| 2 | 6 | 29 | Propionyl-L-Carnitine HCl | 35_36 | 68.1488056 | 4.77683333 | 77.2037114 |
| 2 | 16 | 33 | Pyrimidine | 11_12 | 67.4637517 | 5.1584 | 77.6565191 |
| 2 | 12 | 36 | Pyrimidine | 37_38 | 66.9848642 | 4.659 | 64.9779038 |
| 2 | 15 | 43 | Pyrimidine | 15_16 | | | |
| 2 | 21 | 25 | Pyrimidine | 21_22 | 69.0715419 | 5.0034 | 82.6374037 |

| | | | | | | | |
|---|---|---|---|---|---|---|---|
| 2 | 14 | 48 | Safrole | 13_14 | 67.6913545 | 4.49163333 | 75.8452884 |
| 2 | 7 | 49 | Safrole | 19_20 | 66.6149494 | 5.09626667 | 69.2795768 |
| 2 | 5 | 34 | Safrole | 23_24 | 67.6954215 | 4.6777 | 72.6756345 |
| 2 | 23 | 45 | Safrole | 27_28 | 65.865856 | 4.97683333 | 65.8835192 |
| 3 | 19 | 29 | Control | 33_34 | 66.4109557 | 4.66863333 | 79.8450896 |
| 3 | 23 | 47 | Control | 39_40 | 66.5705092 | 4.76223333 | 71.6945512 |
| 3 | 18 | 25 | Control | 35_36 | | | |
| 3 | 13 | 38 | Control | 17_18 | 75.3652267 | 5.08 | 76.2226281 |
| 3 | 10 | 28 | Dimethylglyoxime | 5_6 | 69.1052849 | 5.1431 | 74.864205 |
| 3 | 16 | 48 | Dimethylglyoxime | 31_32 | 67.6624179 | 4.5546 | 80.750705 |
| 3 | 11 | 45 | Dimethylglyoxime | 29_30 | 68.6402621 | 5.17553333 | 74.864205 |
| 3 | 24 | 49 | Dimethylglyoxime | 15_16 | 66.5361379 | 4.97126667 | 77.5810512 |
| 3 | 4 | 27 | Propionyl-L-Carnitine HCl | 13_14 | 67.7084372 | 5.36106667 | 84.8259743 |
| 3 | 15 | 37 | Propionyl-L-Carnitine HCl | 3_4 | 69.3075621 | 5.0778 | 71.6945512 |
| 3 | 21 | 34 | Propionyl-L-Carnitine HCl | 37_38 | 68.1298129 | 4.92383333 | 73.0529743 |
| 3 | 2 | 31 | Propionyl-L-Carnitine HCl | 21_22 | 67.6406168 | 5.39956667 | 73.0529743 |
| 3 | 20 | 39 | Pyrimidine | 23_24 | 66.1800773 | 4.8939 | 70.7889358 |
| 3 | 3 | 22 | Pyrimidine | 25_26 | 68.6722424 | 5.08033333 | 69.8833204 |
| 3 | 5 | 32 | Pyrimidine | 19_20 | 67.4951588 | 5.02663333 | 84.3731666 |
| 3 | 22 | 44 | Pyrimidine | 11_12 | 69.1177561 | 5.0442 | 76.2226281 |
| 3 | 6 | 41 | Safrole | 9_10 | 67.2734883 | 4.8911 | 72.1473589 |
| 3 | 8 | 36 | Safrole | 27_28 | 67.9034645 | 4.80716667 | 72.6001666 |
| 3 | 14 | 35 | Safrole | 1_2 | 69.3488315 | 5.34986667 | 76.2226281 |
| 3 | 9 | 26 | Safrole | 7_8 | 67.4108383 | 4.91623333 | 83.9203589 |

163

| Run | Bottle | Module | TRT | F57 BagIDs | 48h.gas.prod | CH4.prod | NH3 |
|---|---|---|---|---|---|---|---|
| 1 | 14 | 32 | Control | 29_30 | 107.655029 | 5.34367266 | 26.812 |
| 1 | 24 | 31 | Control | 11_12 | 113.088721 | 5.7251165 | 25.32 |
| 1 | 9 | 47 | Control | 13_14 | 106.749413 | 5.56893898 | 22.76 |
| 1 | 19 | 40 | Control | 15_16 | 111.730298 | 5.54163656 | 24.44 |
| 1 | 13 | 49 | Dimethylglyoxime | 27_28 | 104.485375 | 5.42909491 | 26.078 |
| 1 | 10 | 36 | Dimethylglyoxime | 5_6 | 102.221336 | 5.17260407 | 25.562 |
| 1 | 17 | 38 | Dimethylglyoxime | 39_40 | 112.635913 | 5.97150558 | 26.488 |
| 1 | 22 | 37 | Dimethylglyoxime | 17_18 | 110.371875 | 5.80243342 | 26.054 |
| 1 | 7 | 29 | Propionyl-L-Carnitine HCl | 7_8 | 108.560644 | 5.73051835 | 25.584 |
| 1 | 12 | 44 | Propionyl-L-Carnitine HCl | 1_2 | 111.27749 | 5.38460648 | 25.82 |
| 1 | 1 | 25 | Propionyl-L-Carnitine HCl | 9_10 | 113.994336 | 5.40074767 | 27.35 |
| 1 | 11 | 41 | Propionyl-L-Carnitine HCl | 31_32 | 104.032567 | 5.37616033 | 26.38 |
| 1 | 16 | 34 | Pyrimidine | 21_22 | 104.485375 | 5.31753936 | 23.832 |

| | | | | | | | |
|---|---|---|---|---|---|---|---|
| 1 | 3 | 39 | Pyrimidine | 23_24 | 110.371875 | 5.30520812 | 25.99 |
| 1 | 21 | 45 | Pyrimidine | 3_4 | 104.032567 | 5.40015718 | 25.34 |
| 1 | 6 | 48 | Pyrimidine | 25_26 | 108.560644 | 5.41207379 | 26.532 |
| 1 | 15 | 33 | Safrole | 37_38 | 106.296606 | 5.30600766 | 26.38 |
| 1 | 20 | 35 | Safrole | 19_20 | 111.27749 | 5.21776443 | 21.8 |
| 1 | 4 | 43 | Safrole | 33_34 | 105.39099 | 5.18646628 | 26.326 |
| 1 | 5 | 26 | Safrole | 35_36 | 106.296606 | 5.39224964 | 26.1 |
| 2 | 4 | 39 | Control | 29_30 | 93.8443941 | 4.39786112 | 27.572 |
| 2 | 1 | 41 | Control | 3_4 | 93.84 | 4.80892464 | 28.148 |
| 2 | 10 | 46 | Control | 9_10 | 99.2780864 | 4.99613661 | 27.274 |
| 2 | 18 | 40 | Control | 1_2 | 101.089317 | 5.20020296 | 29.074 |
| 2 | 17 | 37 | Dimethylglyoxime | 31_32 | 95.2028172 | 4.59061638 | 24.484 |
| 2 | 8 | 26 | Dimethylglyoxime | 39_40 | 90.2219326 | 4.22253682 | 27.488 |
| 2 | 13 | 35 | Dimethylglyoxime | 25_26 | 92.0331634 | 4.59049148 | 27.296 |
| 2 | 19 | 32 | Dimethylglyoxime | 5_6 | 93.3915864 | 4.4648027 | 29.51 |
| 2 | 20 | 27 | Propionyl-L-Carnitine HCl | 33_34 | 98.372471 | 5.12379574 | 20.644 |
| 2 | 3 | 47 | Propionyl-L-Carnitine HCl | 7_8 | 91.127548 | 4.12883732 | 28.232 |
| 2 | 24 | 38 | Propionyl-L-Carnitine HCl | 17_18 | 105.164586 | 5.34714598 | 19.018 |
| 2 | 6 | 29 | Propionyl-L-Carnitine HCl | 35_36 | 97.4668557 | 4.65582925 | 27.7 |
| 2 | 16 | 33 | Pyrimidine | 11_12 | 98.8252787 | 5.09780318 | 29 |
| 2 | 12 | 36 | Pyrimidine | 37_38 | 82.5242018 | 3.84480256 | 26.06 |
| 2 | 15 | 43 | Pyrimidine | 15_16 | | | |
| 2 | 21 | 25 | Pyrimidine | 21_22 | 102.44774 | 5.12587024 | 21.046 |
| 2 | 14 | 48 | Safrole | 13_14 | 93.8443941 | 4.21514609 | 28.062 |
| 2 | 7 | 49 | Safrole | 19_20 | 88.41 | 4.50560936 | 27.742 |
| 2 | 5 | 34 | Safrole | 23_24 | 93.3915864 | 4.36857824 | 27.85 |
| 2 | 23 | 45 | Safrole | 27_28 | 86.5994711 | 4.30991134 | 26.73 |
| 3 | 19 | 29 | Control | 33_34 | 103.692961 | 4.84104416 | 24.604 |
| 3 | 23 | 47 | Control | 39_40 | 100.523308 | 4.78715446 | 27.912 |
| 3 | 18 | 25 | Control | 35_36 | | | |
| 3 | 13 | 38 | Control | 17_18 | 100.976115 | 5.12958666 | 20.39 |
| 3 | 10 | 28 | Dimethylglyoxime | 5_6 | 100.523308 | 5.17001423 | 26.814 |
| 3 | 16 | 48 | Dimethylglyoxime | 31_32 | 105.504192 | 4.80529394 | 23.152 |
| 3 | 11 | 45 | Dimethylglyoxime | 29_30 | 98.2592691 | 5.08544123 | 18.912 |
| 3 | 24 | 49 | Dimethylglyoxime | 15_16 | 100.523308 | 4.99728168 | 30.176 |
| 3 | 4 | 27 | Propionyl-L-Carnitine HCl | 13_14 | 110.937885 | 5.94745395 | 25.82 |
| 3 | 15 | 37 | Propionyl-L-Carnitine HCl | 3_4 | 97.8064614 | 4.9664165 | 29.602 |
| 3 | 21 | 34 | Propionyl-L-Carnitine HCl | 37_38 | 96.4480383 | 4.74894066 | 23.866 |
| 3 | 2 | 31 | Propionyl-L-Carnitine HCl | 21_22 | 96.45 | 5.20788205 | 22.272 |
| 3 | 20 | 39 | Pyrimidine | 23_24 | 94.1839999 | 4.60927077 | 27.578 |

| | | | | | | | |
|---|---|---|---|---|---|---|---|
| 3 | 3 | 22 | Pyrimidine | 25_26 | 95.9952307 | 4.8768777 | 27.046 |
| 3 | 5 | 32 | Pyrimidine | 19_20 | 109.126654 | 5.48539675 | 28.462 |
| 3 | 22 | 44 | Pyrimidine | 11_12 | 99.6176922 | 5.02491563 | 29.152 |
| 3 | 6 | 41 | Safrole | 9_10 | 95.0896153 | 4.65092817 | 26.392 |
| 3 | 8 | 36 | Safrole | 27_28 | 95.9952307 | 4.61465073 | 27.702 |
| 3 | 14 | 35 | Safrole | 1_2 | 100.523308 | 5.37786292 | 26.454 |
| 3 | 9 | 26 | Safrole | 7_8 | 107.768231 | 5.29813768 | 25.694 |

| **Run** | **Bottle** | **Module** | **TRT** | **F57 BagIDs** | **Acetate** | **Propionate** | **Isobutyrate** |
|---|---|---|---|---|---|---|---|
| 1 | 14 | 32 | Control | 29_30 | 39.07667 | 18.0029 | 0.774941 |
| 1 | 24 | 31 | Control | 11_12 | 52.68398 | 23.50458 | 1.049111 |
| 1 | 9 | 47 | Control | 13_14 | 51.71933 | 23.61549 | 1.046301 |
| 1 | 19 | 40 | Control | 15_16 | 49.17178 | 22.75816 | 0.990034 |
| 1 | 13 | 49 | Dimethylglyoxime | 27_28 | 50.31458 | 23.08582 | 0.985614 |
| 1 | 10 | 36 | Dimethylglyoxime | 5_6 | 52.10479 | 23.63965 | 1.037949 |
| 1 | 17 | 38 | Dimethylglyoxime | 39_40 | 49.9045 | 22.92057 | 1.001055 |
| 1 | 22 | 37 | Dimethylglyoxime | 17_18 | 51.51058 | 24.03258 | 1.027724 |
| 1 | 7 | 29 | Propionyl-L-Carnitine HCl | 7_8 | 49.43867 | 23.21667 | 0.994957 |
| 1 | 12 | 44 | Propionyl-L-Carnitine HCl | 1_2 | 47.59702 | 21.7904 | 0.94855 |
| 1 | 1 | 25 | Propionyl-L-Carnitine HCl | 9_10 | 50.52471 | 22.80033 | 1.008067 |
| 1 | 11 | 41 | Propionyl-L-Carnitine HCl | 31_32 | 45.78509 | 20.84496 | 0.908966 |
| 1 | 16 | 34 | Pyrimidine | 21_22 | 47.6894 | 21.57107 | 0.961912 |
| 1 | 3 | 39 | Pyrimidine | 23_24 | 53.13426 | 23.5931 | 1.010526 |
| 1 | 21 | 45 | Pyrimidine | 3_4 | 49.10888 | 22.17657 | 0.980905 |
| 1 | 6 | 48 | Pyrimidine | 25_26 | 49.97058 | 23.2541 | 1.015819 |
| 1 | 15 | 33 | Safrole | 37_38 | 50.89195 | 23.45677 | 1.021328 |
| 1 | 20 | 35 | Safrole | 19_20 | 51.35425 | 23.67557 | 1.030055 |
| 1 | 4 | 43 | Safrole | 33_34 | 49.13303 | 22.94148 | 0.983582 |
| 1 | 5 | 26 | Safrole | 35_36 | 50.31738 | 23.56571 | 1.017163 |
| 2 | 4 | 39 | Control | 29_30 | 55.85655 | 25.12772 | 1.090787 |
| 2 | 1 | 41 | Control | 3_4 | 53.27121 | 24.24689 | 1.052667 |
| 2 | 10 | 46 | Control | 9_10 | 57.29577 | 25.93161 | 1.139236 |
| 2 | 18 | 40 | Control | 1_2 | 56.23184 | 25.25505 | 1.114006 |
| 2 | 17 | 37 | Dimethylglyoxime | 31_32 | 54.24964 | 25.15009 | 1.064098 |
| 2 | 8 | 26 | Dimethylglyoxime | 39_40 | 55.64298 | 25.1361 | 1.101364 |
| 2 | 13 | 35 | Dimethylglyoxime | 25_26 | 55.42848 | 25.00338 | 1.089041 |
| 2 | 19 | 32 | Dimethylglyoxime | 5_6 | 55.8614 | 24.90747 | 1.106628 |
| 2 | 20 | 27 | Propionyl-L-Carnitine HCl | 33_34 | 55.21983 | 25.71369 | 1.087928 |
| 2 | 3 | 47 | Propionyl-L-Carnitine HCl | 7_8 | 56.70835 | 25.50852 | 1.108624 |

| | | | | | | | |
|---|---|---|---|---|---|---|---|
| 2 | 24 | 38 | Propionyl-L-Carnitine HCl | 17_18 | 56.8189 | 25.6095 | 1.131686 |
| 2 | 6 | 29 | Propionyl-L-Carnitine HCl | 35_36 | 53.51169 | 24.93651 | 1.059778 |
| 2 | 16 | 33 | Pyrimidine | 11_12 | 54.70216 | 24.97982 | 1.096151 |
| 2 | 12 | 36 | Pyrimidine | 37_38 | 54.9066 | 25.00237 | 1.081651 |
| 2 | 15 | 43 | Pyrimidine | 15_16 | | | |
| 2 | 21 | 25 | Pyrimidine | 21_22 | 55.65192 | 25.02182 | 1.103449 |
| 2 | 14 | 48 | Safrole | 13_14 | 54.86045 | 25.0979 | 1.077896 |
| 2 | 7 | 49 | Safrole | 19_20 | 53.9712 | 23.98457 | 1.053775 |
| 2 | 5 | 34 | Safrole | 23_24 | 56.99309 | 25.80766 | 1.126802 |
| 2 | 23 | 45 | Safrole | 27_28 | 45.53817 | 20.70728 | 0.915354 |
| 3 | 19 | 29 | Control | 33_34 | 53.77419 | 25.86291 | 1.097364 |
| 3 | 23 | 47 | Control | 39_40 | 54.59236 | 26.43021 | 1.100251 |
| 3 | 18 | 25 | Control | 35_36 | | | |
| 3 | 13 | 38 | Control | 17_18 | 55.55609 | 25.8607 | 1.131941 |
| 3 | 10 | 28 | Dimethylglyoxime | 5_6 | 53.9584 | 26.07473 | 1.108928 |
| 3 | 16 | 48 | Dimethylglyoxime | 31_32 | 54.94266 | 25.84041 | 1.128303 |
| 3 | 11 | 45 | Dimethylglyoxime | 29_30 | 53.58905 | 25.15212 | 1.086804 |
| 3 | 24 | 49 | Dimethylglyoxime | 15_16 | 54.33689 | 25.75832 | 1.100547 |
| 3 | 4 | 27 | Propionyl-L-Carnitine HCl | 13_14 | 49.76638 | 25.17727 | 1.077182 |
| 3 | 15 | 37 | Propionyl-L-Carnitine HCl | 3_4 | 54.09707 | 25.45392 | 1.106074 |
| 3 | 21 | 34 | Propionyl-L-Carnitine HCl | 37_38 | 53.76715 | 24.80978 | 1.083153 |
| 3 | 2 | 31 | Propionyl-L-Carnitine HCl | 21_22 | 47.50706 | 23.28313 | 1.046941 |
| 3 | 20 | 39 | Pyrimidine | 23_24 | 54.59276 | 25.72412 | 1.096366 |
| 3 | 3 | 22 | Pyrimidine | 25_26 | 49.5949 | 24.95782 | 1.069893 |
| 3 | 5 | 32 | Pyrimidine | 19_20 | 51.61553 | 25.06978 | 1.09821 |
| 3 | 22 | 44 | Pyrimidine | 11_12 | 54.85836 | 25.00725 | 1.12657 |
| 3 | 6 | 41 | Safrole | 9_10 | 50.74868 | 24.5603 | 1.065885 |
| 3 | 8 | 36 | Safrole | 27_28 | 53.66077 | 25.33746 | 1.113016 |
| 3 | 14 | 35 | Safrole | 1_2 | 54.75607 | 25.44825 | 1.12757 |
| 3 | 9 | 26 | Safrole | 7_8 | 53.48065 | 26.00589 | 1.100633 |

| Run | Bottle | Module | TRT | F57 BagIDs | Butyrate | Isovalerate | Valerate |
|---|---|---|---|---|---|---|---|
| 1 | 14 | 32 | Control | 29_30 | 10.68307 | 1.916212 | 1.9435 |
| 1 | 24 | 31 | Control | 11_12 | 13.98744 | 2.560564 | 2.46084 |
| 1 | 9 | 47 | Control | 13_14 | 13.82888 | 2.547251 | 2.480319 |
| 1 | 19 | 40 | Control | 15_16 | 13.05293 | 2.404567 | 2.433913 |
| 1 | 13 | 49 | Dimethylglyoxime | 27_28 | 13.41149 | 2.416514 | 2.453951 |
| 1 | 10 | 36 | Dimethylglyoxime | 5_6 | 13.55685 | 2.493915 | 2.492389 |
| 1 | 17 | 38 | Dimethylglyoxime | 39_40 | 13.47854 | 2.449082 | 2.44277 |

| | | | | | | | |
|---|---|---|---|---|---|---|---|
| 1 | 22 | 37 | Dimethylglyoxime | 17_18 | 13.65014 | 2.504059 | 2.476292 |
| 1 | 7 | 29 | Propionyl-L-Carnitine HCl | 7_8 | 13.10628 | 2.453178 | 2.458969 |
| 1 | 12 | 44 | Propionyl-L-Carnitine HCl | 1_2 | 12.6856 | 2.310297 | 2.323231 |
| 1 | 1 | 25 | Propionyl-L-Carnitine HCl | 9_10 | 13.14224 | 2.414307 | 2.422698 |
| 1 | 11 | 41 | Propionyl-L-Carnitine HCl | 31_32 | 12.60373 | 2.283594 | 2.311712 |
| 1 | 16 | 34 | Pyrimidine | 21_22 | 13.06558 | 2.346153 | 2.315386 |
| 1 | 3 | 39 | Pyrimidine | 23_24 | 13.67566 | 2.455552 | 2.510357 |
| 1 | 21 | 45 | Pyrimidine | 3_4 | 12.98921 | 2.383449 | 2.346819 |
| 1 | 6 | 48 | Pyrimidine | 25_26 | 13.63696 | 2.505428 | 2.535695 |
| 1 | 15 | 33 | Safrole | 37_38 | 13.57262 | 2.488351 | 4.021847 |
| 1 | 20 | 35 | Safrole | 19_20 | 13.5687 | 2.518452 | 2.469349 |
| 1 | 4 | 43 | Safrole | 33_34 | 13.18461 | 2.427102 | 2.507241 |
| 1 | 5 | 26 | Safrole | 35_36 | 13.70263 | 2.471212 | 2.496753 |
| 2 | 4 | 39 | Control | 29_30 | 16.47803 | 3.461312 | 3.573021 |
| 2 | 1 | 41 | Control | 3_4 | 15.48569 | 3.3263 | 3.571895 |
| 2 | 10 | 46 | Control | 9_10 | 16.55893 | 3.488869 | 3.389195 |
| 2 | 18 | 40 | Control | 1_2 | 16.21022 | 3.530408 | 3.404101 |
| 2 | 17 | 37 | Dimethylglyoxime | 31_32 | 15.74472 | 3.35086 | 3.412314 |
| 2 | 8 | 26 | Dimethylglyoxime | 39_40 | 16.20394 | 3.365356 | 3.479778 |
| 2 | 13 | 35 | Dimethylglyoxime | 25_26 | 16.19394 | 3.366271 | 3.464133 |
| 2 | 19 | 32 | Dimethylglyoxime | 5_6 | 16.15319 | 3.465684 | 3.487815 |
| 2 | 20 | 27 | Propionyl-L-Carnitine HCl | 33_34 | 16.16755 | 3.37152 | 3.409607 |
| 2 | 3 | 47 | Propionyl-L-Carnitine HCl | 7_8 | 16.32336 | 3.471796 | 3.462474 |
| 2 | 24 | 38 | Propionyl-L-Carnitine HCl | 17_18 | 16.63796 | 3.516014 | 3.348692 |
| 2 | 6 | 29 | Propionyl-L-Carnitine HCl | 35_36 | 15.71824 | 3.308436 | 3.431738 |
| 2 | 16 | 33 | Pyrimidine | 11_12 | 15.90128 | 3.347656 | 3.338585 |
| 2 | 12 | 36 | Pyrimidine | 37_38 | 16.20955 | 3.356356 | 3.419682 |
| 2 | 15 | 43 | Pyrimidine | 15_16 | | | |
| 2 | 21 | 25 | Pyrimidine | 21_22 | 16.33845 | 3.463133 | 3.320795 |
| 2 | 14 | 48 | Safrole | 13_14 | 15.65267 | 3.347308 | 4.447622 |
| 2 | 7 | 49 | Safrole | 19_20 | 15.83968 | 3.336268 | 3.295992 |
| 2 | 5 | 34 | Safrole | 23_24 | 16.65481 | 3.506929 | 3.565679 |
| 2 | 23 | 45 | Safrole | 27_28 | 13.35674 | 2.94413 | 2.958642 |
| 3 | 19 | 29 | Control | 33_34 | 13.9044 | 2.892004 | 2.446449 |
| 3 | 23 | 47 | Control | 39_40 | 13.98152 | 2.951699 | 2.418678 |
| 3 | 18 | 25 | Control | 35_36 | | | |
| 3 | 13 | 38 | Control | 17_18 | 14.32485 | 3.046863 | 2.457747 |
| 3 | 10 | 28 | Dimethylglyoxime | 5_6 | 14.2825 | 2.980141 | 2.4703 |
| 3 | 16 | 48 | Dimethylglyoxime | 31_32 | 13.98897 | 3.001548 | 2.510366 |
| 3 | 11 | 45 | Dimethylglyoxime | 29_30 | 14.05413 | 2.84368 | 2.398213 |

| | | | | | | | |
|---|---|---|---|---|---|---|---|
| 3 | 24 | 49 | Dimethylglyoxime | 15_16 | 14.15286 | 2.916545 | 2.435474 |
| 3 | 4 | 27 | Propionyl-L-Carnitine HCl | 13_14 | 13.73946 | 2.844176 | 2.376271 |
| 3 | 15 | 37 | Propionyl-L-Carnitine HCl | 3_4 | 14.14907 | 2.984307 | 2.430076 |
| 3 | 21 | 34 | Propionyl-L-Carnitine HCl | 37_38 | 13.83345 | 2.954473 | 2.397395 |
| 3 | 2 | 31 | Propionyl-L-Carnitine HCl | 21_22 | 13.11039 | 2.806713 | 2.246622 |
| 3 | 20 | 39 | Pyrimidine | 23_24 | 14.05344 | 2.989503 | 2.428439 |
| 3 | 3 | 22 | Pyrimidine | 25_26 | 13.68043 | 2.893125 | 2.385328 |
| 3 | 5 | 32 | Pyrimidine | 19_20 | 13.88574 | 2.938221 | 2.419839 |
| 3 | 22 | 44 | Pyrimidine | 11_12 | 13.97263 | 3.033439 | 2.456693 |
| 3 | 6 | 41 | Safrole | 9_10 | 13.19263 | 2.607588 | 3.963031 |
| 3 | 8 | 36 | Safrole | 27_28 | 13.99659 | 2.992919 | 2.438089 |
| 3 | 14 | 35 | Safrole | 1_2 | 14.42752 | 3.017506 | 2.440047 |
| 3 | 9 | 26 | Safrole | 7_8 | 13.91864 | 2.937214 | 2.458037 |

166

| Run | Bottle | Module | TRT | F57 BagIDs | Total | A:P |
|---|---|---|---|---|---|---|
| 1 | 14 | 32 | Control | 29_30 | 72.3973 | 2.170576 |
| 1 | 24 | 31 | Control | 11_12 | 96.24651 | 2.241435 |
| 1 | 9 | 47 | Control | 13_14 | 95.23757 | 2.190059 |
| 1 | 19 | 40 | Control | 15_16 | 90.81139 | 2.160622 |
| 1 | 13 | 49 | Dimethylglyoxime | 27_28 | 92.66797 | 2.179458 |
| 1 | 10 | 36 | Dimethylglyoxime | 5_6 | 95.32555 | 2.204127 |
| 1 | 17 | 38 | Dimethylglyoxime | 39_40 | 92.19652 | 2.17728 |
| 1 | 22 | 37 | Dimethylglyoxime | 17_18 | 95.20138 | 2.143364 |
| 1 | 7 | 29 | Propionyl-L-Carnitine HCl | 7_8 | 91.66873 | 2.129447 |
| 1 | 12 | 44 | Propionyl-L-Carnitine HCl | 1_2 | 87.6551 | 2.184311 |
| 1 | 1 | 25 | Propionyl-L-Carnitine HCl | 9_10 | 92.31236 | 2.215965 |
| 1 | 11 | 41 | Propionyl-L-Carnitine HCl | 31_32 | 84.73805 | 2.196458 |
| 1 | 16 | 34 | Pyrimidine | 21_22 | 87.9495 | 2.210803 |
| 1 | 3 | 39 | Pyrimidine | 23_24 | 96.37946 | 2.25211 |
| 1 | 21 | 45 | Pyrimidine | 3_4 | 89.98583 | 2.214449 |
| 1 | 6 | 48 | Pyrimidine | 25_26 | 92.91859 | 2.148893 |
| 1 | 15 | 33 | Safrole | 37_38 | 95.45287 | 2.169606 |
| 1 | 20 | 35 | Safrole | 19_20 | 94.61637 | 2.169082 |
| 1 | 4 | 43 | Safrole | 33_34 | 91.17704 | 2.141668 |
| 1 | 5 | 26 | Safrole | 35_36 | 93.57084 | 2.135195 |
| 2 | 4 | 39 | Control | 29_30 | 105.5874 | 2.222906 |
| 2 | 1 | 41 | Control | 3_4 | 100.9547 | 2.197032 |
| 2 | 10 | 46 | Control | 9_10 | 107.8036 | 2.209495 |
| 2 | 18 | 40 | Control | 1_2 | 105.7456 | 2.226558 |

| | | | | | | |
|---|---|---|---|---|---|---|
| 2 | 17 | 37 | Dimethylglyoxime | 31_32 | 102.9717 | 2.157035 |
| 2 | 8 | 26 | Dimethylglyoxime | 39_40 | 104.9295 | 2.213668 |
| 2 | 13 | 35 | Dimethylglyoxime | 25_26 | 104.5452 | 2.21684 |
| 2 | 19 | 32 | Dimethylglyoxime | 5_6 | 104.9822 | 2.242757 |
| 2 | 20 | 27 | Propionyl-L-Carnitine HCl | 33_34 | 104.9701 | 2.147488 |
| 2 | 3 | 47 | Propionyl-L-Carnitine HCl | 7_8 | 106.5831 | 2.223114 |
| 2 | 24 | 38 | Propionyl-L-Carnitine HCl | 17_18 | 107.0627 | 2.218665 |
| 2 | 6 | 29 | Propionyl-L-Carnitine HCl | 35_36 | 101.9664 | 2.145917 |
| 2 | 16 | 33 | Pyrimidine | 11_12 | 103.3657 | 2.189854 |
| 2 | 12 | 36 | Pyrimidine | 37_38 | 103.9762 | 2.196056 |
| 2 | 15 | 43 | Pyrimidine | 15_16 | | |
| 2 | 21 | 25 | Pyrimidine | 21_22 | 104.8996 | 2.224135 |
| 2 | 14 | 48 | Safrole | 13_14 | 104.4838 | 2.185858 |
| 2 | 7 | 49 | Safrole | 19_20 | 101.4815 | 2.250247 |
| 2 | 5 | 34 | Safrole | 23_24 | 107.655 | 2.208379 |
| 2 | 23 | 45 | Safrole | 27_28 | 86.42032 | 2.199138 |
| 3 | 19 | 29 | Control | 33_34 | 99.97731 | 2.079201 |
| 3 | 23 | 47 | Control | 39_40 | 101.4747 | 2.065529 |
| 3 | 18 | 25 | Control | 35_36 | | |
| 3 | 13 | 38 | Control | 17_18 | 102.3782 | 2.148282 |
| 3 | 10 | 28 | Dimethylglyoxime | 5_6 | 100.875 | 2.069375 |
| 3 | 16 | 48 | Dimethylglyoxime | 31_32 | 101.4123 | 2.12623 |
| 3 | 11 | 45 | Dimethylglyoxime | 29_30 | 99.124 | 2.130598 |
| 3 | 24 | 49 | Dimethylglyoxime | 15_16 | 100.7006 | 2.109488 |
| 3 | 4 | 27 | Propionyl-L-Carnitine HCl | 13_14 | 94.98074 | 1.97664 |
| 3 | 15 | 37 | Propionyl-L-Carnitine HCl | 3_4 | 100.2205 | 2.125294 |
| 3 | 21 | 34 | Propionyl-L-Carnitine HCl | 37_38 | 98.84541 | 2.167176 |
| 3 | 2 | 31 | Propionyl-L-Carnitine HCl | 21_22 | 90.00087 | 2.040407 |
| 3 | 20 | 39 | Pyrimidine | 23_24 | 100.8846 | 2.12224 |
| 3 | 3 | 22 | Pyrimidine | 25_26 | 94.5815 | 1.987149 |
| 3 | 5 | 32 | Pyrimidine | 19_20 | 97.02731 | 2.058874 |
| 3 | 22 | 44 | Pyrimidine | 11_12 | 100.4549 | 2.193698 |
| 3 | 6 | 41 | Safrole | 9_10 | 96.13812 | 2.066289 |
| 3 | 8 | 36 | Safrole | 27_28 | 99.53884 | 2.117843 |
| 3 | 14 | 35 | Safrole | 1_2 | 101.217 | 2.151663 |
| 3 | 9 | 26 | Safrole | 7_8 | 99.90106 | 2.056482 |
